# Multi-chamber cardioids unravel human heart development and cardiac defects

**DOI:** 10.1101/2022.07.14.499699

**Authors:** Clara Schmidt, Alison Deyett, Tobias Ilmer, Aranxa Torres Caballero, Simon Haendeler, Lokesh Pimpale, Michael A. Netzer, Lavinia Ceci Ginistrelli, Martina Cirigliano, Estela Juncosa Mancheno, Daniel Reumann, Katherina Tavernini, Steffen Hering, Pablo Hofbauer, Sasha Mendjan

## Abstract

The number one cause of human fetal death are defects in heart development. Because the human embryonic heart is inaccessible, and the impacts of mutations, drugs, and environmental factors on the specialized functions of different heart compartments are not captured by *in vitro* models, determining the underlying causes is difficult. Here, we established a human cardioid platform that recapitulates the development of all major embryonic heart compartments, including right and left ventricles, atria, outflow tract, and atrioventricular canal. By leveraging both 2D and 3D differentiation, we efficiently generated progenitor subsets with distinct first, anterior, and posterior second heart field identities. This advance enabled the reproducible generation of cardioids with compartment-specific *in vivo*-like gene expression profiles, morphologies, and functions. We used this platform to unravel the ontogeny of signal and contraction propagation between interacting heart chambers and dissect how genetic and environmental factors cause region-specific defects in the developing human heart.

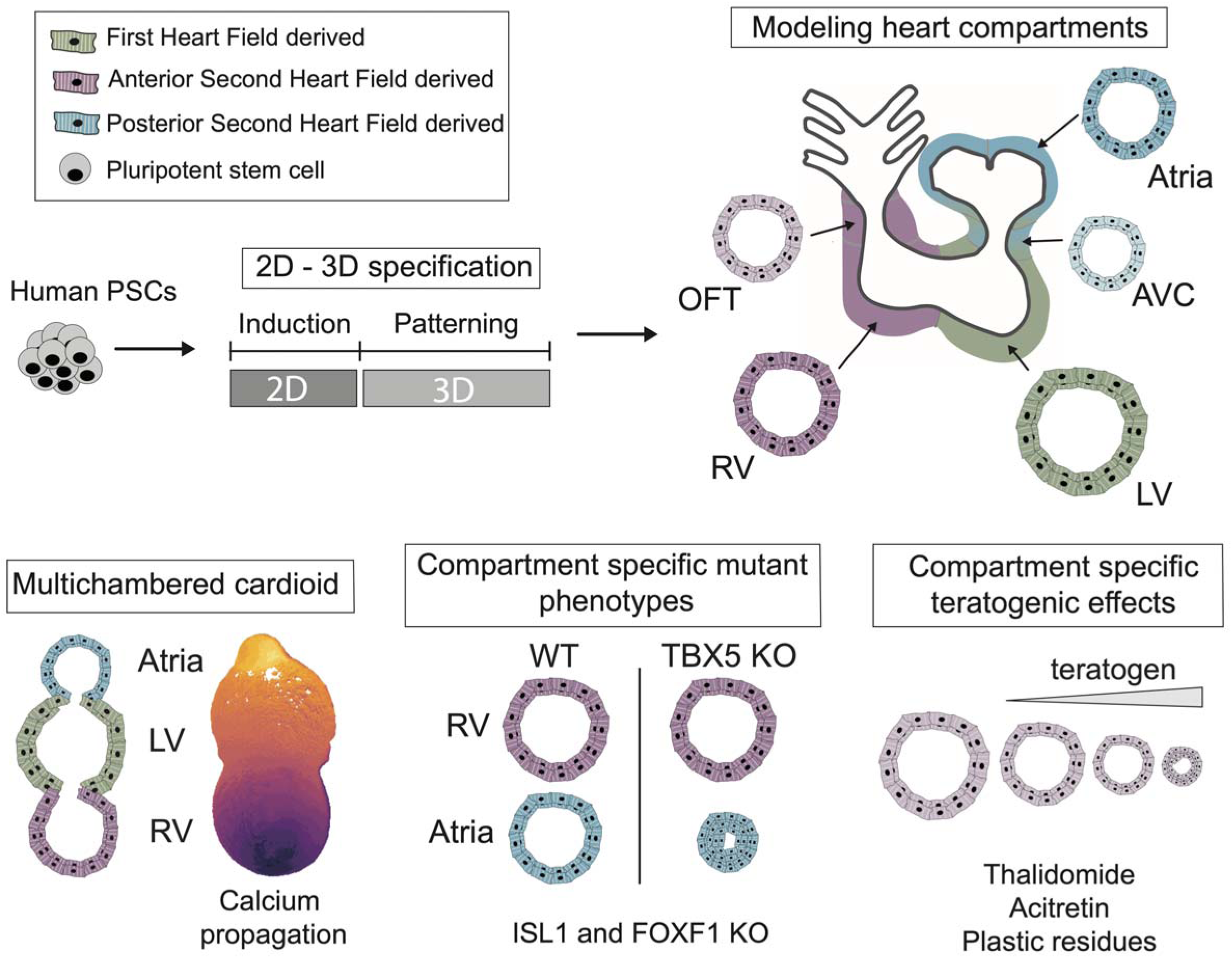

**HIGHLIGHTS:**

- Mesoderm induction and patterning signals specify aSHF, pSHF, and FHF progenitors
- Cardiac progenitors sort, co-develop and functionally connect in multi-chamber cardioids
- Multi-chamber cardioids coordinate contraction propagation and share a lumen
- Multi-chamber platform dissects genetic (ISL1, TBX5, FOXF1) and teratogenic defects

## INTRODUCTION

Congenital heart disease (CHD) is the most common human developmental birth defect and the most prevalent cause of embryonic and fetal mortality (van der Linde et al., 2011; Jin et al., 2017). CHDs most often affect specific compartments of the embryonic heart, such as the outflow tract (OFT), the atria, the atrioventricular canal (AVC), and the right ventricle (RV) (Fahed et al., 2013). For about 56% of diagnosed CHD cases, the underlying cause is unknown but is assumed to originate from undiscovered genetic mutations, environmental factors, or a combination of both (Zaidi and Brueckner, 2017). To identify possible causes and preventive measures, we need models encompassing all compartments of the developing human heart.

CHDs occur early in embryonic development, often before pregnancy has been detected, making the characterization of disease etiology particularly challenging (Gonzalez-Teran et al., 2022; Kathiresan and Srivastava, 2012). These difficulties are compounded by the lack of control over the interactions between genetic background and environmental factors during human embryonic development (Zaidi and Brueckner, 2017). Understanding the etiology of CHD solely through animal models is not feasible, given tissue complexity, developmental speed, inaccessibility, and species-specific physiological differences (Srivastava, 2021). Therefore, recently reported human self-organizing cardiac organoid models are important and complementary, as these represent experimental models of human cardiac development and thereby allow reductionist dissection of mechanisms in high-throughput, obtaining results with high statistical significance (Hofbauer et al., 2021a; Kim et al., 2022). However, these systems do not yet allow the mechanistic interrogation of defects representing all interacting compartments (OFT, AVC, atria, RV, and left ventricle (LV)) of the human embryonic heart.

For a controlled *in vitro* system to mimic human heart development, it is essential to deploy the *in vivo* principles that govern the coalescence of all lineages in building a heart (Kelly et al., 2014; Meilhac and Buckingham, 2018). Heart structures are predominantly derived from three progenitor populations that each give rise to specific cardiomyocyte (CM) lineages. The first heart field (FHF) progenitors primarily give rise to the developing LV, the anterior second heart field (aSHF) progenitors give rise to the developing RV and most of the OFT, and the posterior second heart field (pSHF) progenitors give rise to most of the atria and a portion of the AVC. The development of these structures is carefully timed such that the FHF-derived CMs form the heart tube, which gives rise to the LV, while the aSHF and pSHF progenitors differentiate to form the remaining compartments in a delayed and gradual fashion. This complex and dynamic process is orchestrated by developmental signaling through multiple pathways (WNT, Activin/Nodal, BMP, RA, FGF, NOTCH, etc.) at specific stages of early cardiogenesis (Bruneau, 2013). The signaling pathways control key downstream compartment-specific transcription factors (TF) (e. g., TBX1/5, FOXF1, HAND1/2, TBX2/3, ISL1, IRX4, HEY1/2, and NR2F1/2), instructing progenitor specification, morphogenesis, and physiology at specific stages (Christoffels and Jensen, 2020). Although the core components and their roles are well known, we lack a human model enabling mechanistic dissection of how this network becomes perturbed by mutations or environmental factors that lead to CHD or fetal death.

Here, we established a multi-chamber cardioid platform in which all the major compartments of the human embryonic heart are represented. We then use this platform to unravel how mutations and environmental factors impact specific regions of the developing heart and how chambers interact to develop a coordinated sequential contraction pattern.

## RESULTS

### Generation of cardioids from aSHF and pSHF progenitors

To derive progenitor populations from the SHF lineages, we first hypothesized that the aSHF is likely exposed to WNT and TGF-beta signaling inhibition, a similar signaling environment as other anterior and dorsal embryonic regions (neuroectoderm and head mesoderm) (Arkell and Tam, 2012; Nandkishore et al., 2018). Thus, we set out to develop a method to derive cardioids from the aSHF lineage by applying these signaling conditions in the 3D cardioid approach (Hofbauer et al., 2021b). The first stage of differentiation consisted of mesoderm induction, followed by the aSHF patterning stage 1 using dual WNT and TGF-beta signaling inhibition (Figure 1A). After 3.5 days of 3D differentiation, we observed a heterogeneous progenitor population in which one subpopulation expressed the aSHF markers TBX1 and FOXC2, and a second subpopulation expressed the FHF and pSHF marker TBX5 (Figures 1B, 1C, and S1A). To determine the origin of this heterogeneity, we analyzed the earlier mesoderm induction stage (day 1.5) by immunostaining and found that the mesoderm marker EOMES was only expressed on the outside of the cardioid. In contrast, the cardioid core still expressed the pluripotency and neuroectoderm marker SOX2 (Figure S1B). Based on these data, we hypothesized that cells in 2D receive more equally distributed mesoderm induction signals, resulting in a homogenous exit from pluripotency and differentiation, whereas mesoderm is not induced homogenously in 3D. When we induced mesoderm in 2D and initiated differentiation in 3D only at the patterning stage 1 (day 1.5), cells exited pluripotency efficiently (Figure S1B), expressed high levels of TBX1 and FOXC2 (67%, protein level) while only a few cells expressed TBX5 (Figures 1C, S1A, S1D, 1G, and 1G’’). In addition, we observed no expression of head mesoderm markers (Bothe et al., 2011) at day 3.5 in aSHF progenitors (Figure S1C), further indicating that the staged 2D-3D differentiation approach produces more homogenous progenitor populations for both FHF and aSHF progenitor cells.

**Figure 1:**
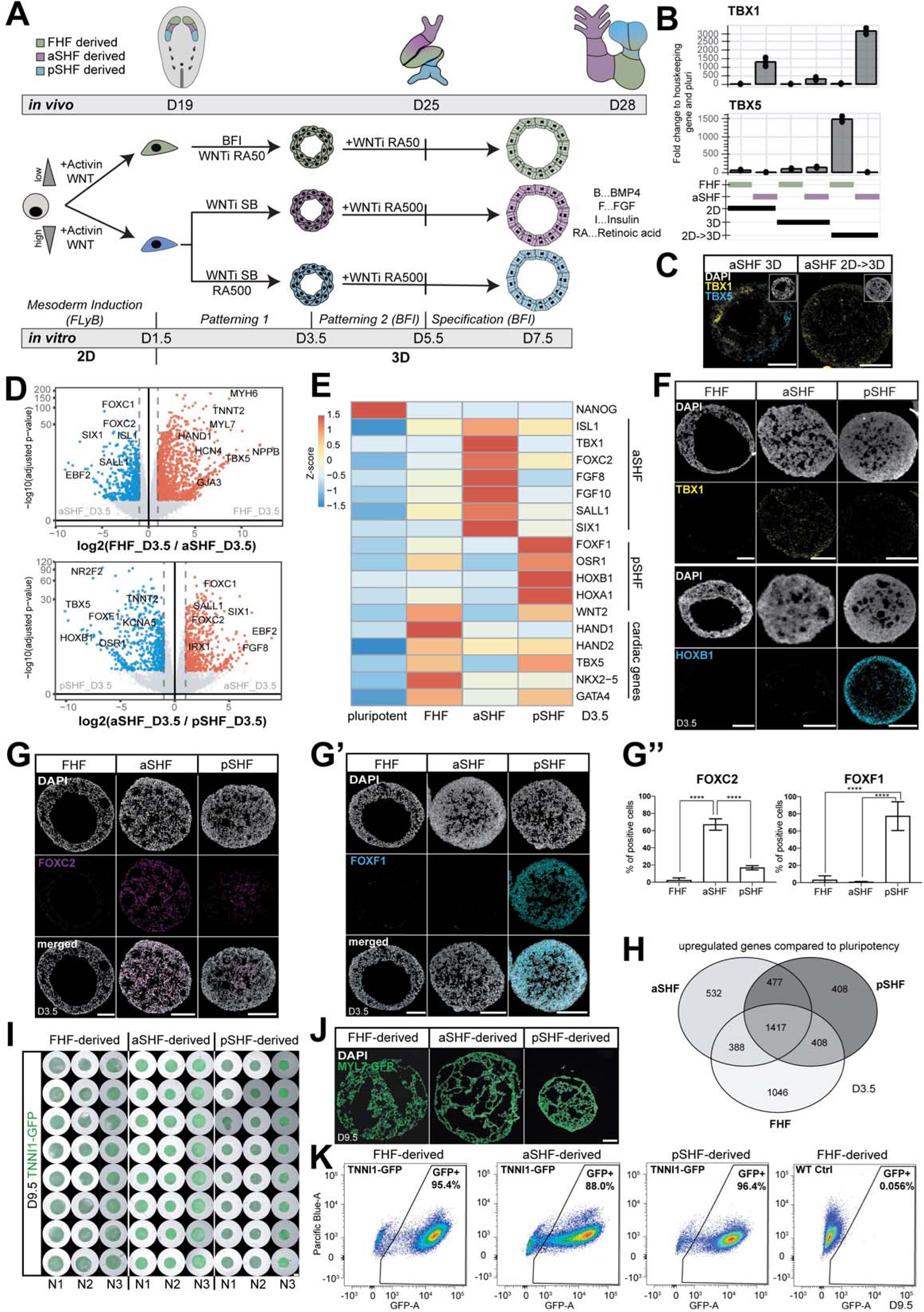
aSHF and pSHF progenitors express specific markers and form functional cardioids. (A) Differentiation protocol into three main cardiac lineages: first heart field (FHF), anterior second heart field (aSHF), and posterior second heart field (pSHF). (B) RT-qPCR of TBX1 and TBX5 levels of FHF and aSHF progenitors at day 3.5 in 2D, 3D, and 2D->3D protocols. Fold change normalized to a housekeeping gene (PBGD) and pluripotency. (C) RNA-scope staining of aSHF cardioid cryosections at day 3.5 for TBX1 and TBX5 expression in 2D->3D approach compared to 3D differentiation. (D) Volcano plot of differentially expressed genes by bulk RNA-seq at day 3.5 of FHF versus aSHF progenitors (top) and aSHF versus pSHF progenitors (bottom). (E) Heatmap of bulk RNA-seq for cardiac mesoderm TFs at day 3.5 for all progenitor populations. (F) RNA-scope staining (TBX1 and HOXB1) of FHF, aSHF, and FHF progenitors at day 3.5. (G) Immunostaining of aSHF marker (FOXC2) and pSHF marker (FOXF1) (G’) of cardioids at day 3.5 and (G’’) quantification of staining (N=3, n=3-4). (H) Venn Diagram showing the number of shared upregulated genes compared to pluripotency between aSHF, pSHF, and FHF at day 3.5. (I) Representative whole-mount images of cardioids derived from TNNI1-GFP reporter line for FHF, aSHF, and pSHF- derived cardioids at day 9.5 across different biological and technical replicates. scale bar 500 µm (J) Cryosection of FHF, aSHF, and pSHF cardioids at day 9.5 tagged for CM-specific marker MYL7. (K) Representative flow cytometry plot of FHF, aSHF and pSHF derived CM using TNNI1-GFP reporter and WT line at day 9.5. All scale bars in this figure have a length of 200µm except where specified. Used cell lines in this figure: H9 and WTC. All bar graphs show mean ± SD. For all statistics, one-way ANOVA was used. *p < 0.05, **p < 0.01, *** p < 0.001, ****p < 0.0001. ns: not significant.

In contrast to the aSHF, the pSHF is exposed to retinoic acid (RA) signaling *in vivo* (Ghyselinck and Duester, 2019), which activates the pSHF regulators (HOXB1, HOXA1, TBX5) and inhibits the aSHF expression signature (TBX1, FOXC2, SIX1). Consistently, we observed that adding RA during the aSHF patterning stage 1 led to a switch toward the pSHF identity (Figures 1A, 1F, 1G’, 1G’’ and S1E). In contrast, manipulation of other signaling pathways (SHH, WNT, FGF, and BMP) had little to no effect on the expression of major aSHF or pSHF markers (Figures S1E and S1F). In line with *in vivo* signaling, different Activin and WNT signaling levels during mesoderm induction promoted the aSHF and pSHF over the FHF lineage (Figure S1G) (REF). Strikingly, when we analyzed the three progenitor subtypes by RNA-seq, we found that the FHF, aSHF, and pSHF primary markers were among the most differentially expressed genes (Figures 1D and E). When compared to the pluripotent state, 532 genes were specifically upregulated in aSHF, 408 in pSHF, and 1046 in FHF cells, and 1417 genes were shared between the three protocols (Figure 1H). Notably, key aSHF markers (TBX1, SIX1, FOXC2) were lowly expressed in FHF and pSHF, while pSHF markers (HOXB1, HOXA1, FOXF1 (77%, protein level)) were hardly detectable in the aSHF and FHF progenitors (Figures 1D-G"). The specificity and homogeneity of the progenitor populations were further underscored by the mutually exclusive expression of lineage-specific markers TBX1 and HOXB1, as seen by RNA hybridization (Figure 1F) and immunostaining for FOXF1, TBX5, and FOXC2 (Figures 1G-G’’ and S1H). Still, all populations were positive for the cardiac progenitor marker NKX2-5 and mostly negative for the pluripotency and neuroectoderm marker SOX2 (Figure S1I). Overall, these data support that by day 3.5 of differentiation in the cardioid system, we can efficiently and homogeneously generate all three major cardiac progenitors.

The FHF, aSHF, and pSHF progenitors give rise to several different cardiac cell types in the embryo, including CMs and endocardial cells. We showed previously that FHF progenitors generate LV chamber-like contracting cardioids (LV cardioids), which contain CM and endocardial-like cells (Hofbauer et al., 2021b). Following this method, we continued to inhibit WNT signaling while treating the a/pSHF progenitors with BMP, FGF, Insulin, and RA (cardiac patterning 2) (Figure 1A), which resulted in the reproducible formation of contracting cavity-containing cardioids in high throughput (N3, n24, for each) (Figures 1I and 1J). In contrast to FHF-derived cardioids, the a/pSHF-derived cardioids required a higher RA dosage at this stage. Efficient aSHF differentiation also required a lower seeding density (Figure S1J), as a high density led to inefficient CM differentiation and expression of neural markers within the organoid core (Figures S1J’). More than 85% of the cardioid cells expressed the key CM marker TNNI1 (Figures 1K and S1L) and lacked expression of endoderm (FOXA2), neuroectoderm (SOX2), and fibroblast (COL1A1) markers (Figure S1K). Finally, aSHF and pSHF cells also differentiated efficiently into PECAM1+ endothelial cells in 2D when exposed to VEGF and Forskolin after the a/pSHF patterning stage 1 (Figure S1M). In summary, by applying *in vivo*-like signaling, a/pSHF progenitors can be differentiated into CM and endothelial lineages within the cardioid system.

### Formation of RV and atrial cardioids

During early cardiac development, FHF progenitors differentiate into CMs that form the heart tube, while aSHF progenitors first proliferate and then differentiate together with pSHF progenitors at a later developmental time (Kelly et al., 2014). Thus, we hypothesized that, relative to FHF-derived cardioids, the proliferation rate would be higher in SHF-derived cardioids, while morphogenesis and differentiation are delayed. A detailed time-course analysis revealed a CM specification and morphogenesis delay in SHF cardioids (Figures 2A-C), while proliferation rate and Ki67 expression elevated in aSHF progenitors until day 4.5 (Figures 2C and 2D). In addition, aSHF progenitors appeared more epithelial-like, as seen by higher CDH1 and lower CDH2 expression (Figure S2A), reminiscent of *in vivo* (Cortes et al., 2018). The a/pSHF cardioids were also smaller than FHF cardioids and started to express the CM marker TNNI1 one day later (Figures 2A and 2B). Global gene expression confirmed that expression of sarcomeric and structural CM genes was delayed in SHF-derived cardioids (Figure 2E and 2F), and we observed a delay in cavity formation (Figure 2C, white and yellow arrows). Taken together, the staggered differentiation of SHF and FHF-derived cardioids *in vitro* is consistent with the *in vivo* developmental timing of SHF and FHF differentiation and morphogenesis. Next, we asked whether the acquisition of chamber identity also followed the developmental trajectory on a/pSHF-derived cardioids. *In vivo*, the FHF gives rise to the LV and a minor portion of atrial CMs, whereas the aSHF and pSHF give rise to the RV and atria, respectively (Meilhac and Buckingham, 2018). To answer that question, we compared the specification potential of a/pSHF and FHF progenitors by adjusting the concentration of RA. We observed that aSHF progenitors gave rise to early RV-like identity (IRX1, IRX2, IRX3, NPPA), while the pSHF progenitors differentiated into early atrial CMs (HEY1, NR2F1, NR2F2) (Figure 2F). In a global gene expression comparison at day 9.5 of FHF/aSHF/pSHF-derived cardioids, we found that the top upregulated genes in aSHF-derived cardioids included ISL1, IRX1, HEY2, and RFTN1 (Figure 2G and S2B), which have been all implicated in ventricular identity and physiology. Moreover, when we compared FHF and aSHF-derived cardioids to pSHF-derived cardioids, we found that TBX5, NR2F2, and NR2F1 were upregulated, consistent with early atrial identity (Figure 2G and S2B). Importantly, these results were confirmed on a protein level for IRX1, NR2F2, and HEY2 (Figures 2H, 2H’, and S2C). When compared to the pluripotent state, 376 genes were specifically upregulated in aSHF-derived, 645 in pSHF-derived, and 449 in FHF-derived cells, and 3508 genes were shared between the three protocols (Figure S2D). The specification of the CM sublineages was also achieved using hESC H9s (Figure S2E). In summary, aSHF progenitors specify into RV-like cardioids (RV cardioids), and pSHF progenitors form atrial cardioids, showing that the early priming of progenitors is crucial to obtaining different chamber identities in the developing heart.

**Figure 2:**
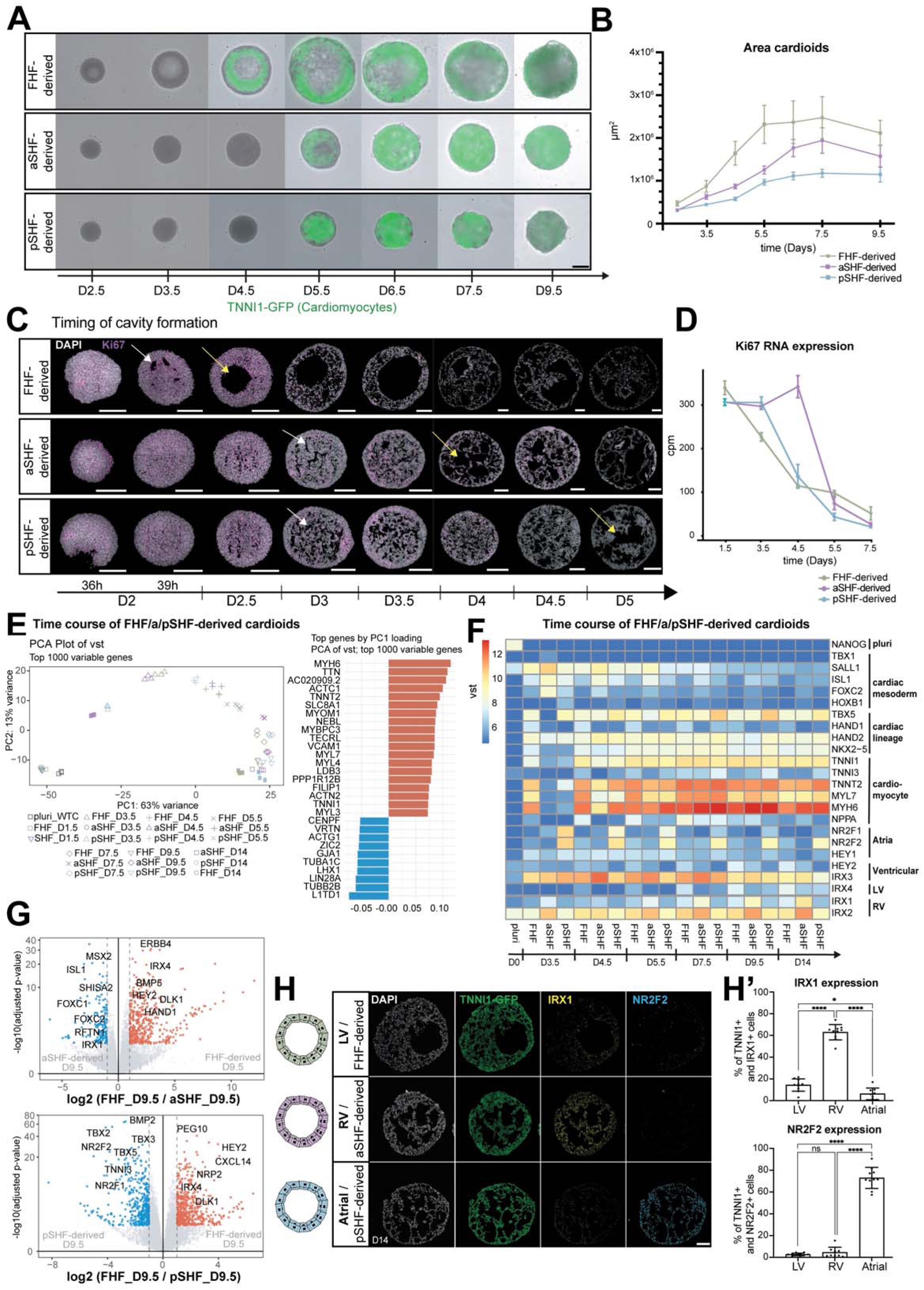
a/pSHF-derived cardioids exhibit *in vivo*-like morphogenesis delay and gene expression. (A) Time course of the three cardioid subtypes from day 2.5 until day 9.5; representative whole-mount images of cardioids derived from the TNNI1-GFP reporter line. Scale bar, 500µm (B) Quantification of cardioid area change during differentiation in A (N=3, n=32). (C) Cryosections of cardioids from day 2 until day 5.5, showing delayed cavity initiation (white arrow) and cavity formation (yellow arrow) in SHF lineages. (C) Immunostaining for Ki67 and (D) mRNA expression by bulk RNA-seq quantification of proliferation marker Ki67 over time (N=3, n=8). Each dot represents the mean ± SD. cpm: counts per million. (E) Principal component analysis (PCA) plot of vst using the top 1000 variable genes. (F) Expression of lineage-specific genes over time shown by bulk RNA-seq. (G) The volcano plot shows the differentially expressed genes at day 9.5 of FHF versus aSHF (top) and FHF versus pSHF (bottom) cardioids. (H) Lineage-specific ICC staining of IRX1 (RV marker) and NR2F2 (atrial marker) at day 14 and (H’) quantification of staining. Each data point represents one cardioid (N=3, n=3-4). All scale bars in this figure have a length of 200µm except where specified. All bar graphs show mean ± SD. For all statistics, one-way ANOVA was used. vst: variance-stabilized transformed counts. *p < 0.05, **p < 0.01, *** p < 0.001, ****p < 0.0001. ns: not significant.

### Specification into OFT, AVC, and chamber cardioids

Besides the RV, aSHF progenitors differentiate into the OFT, which gives rise to the aortic and pulmonary valve and vessel structures (Kelly et al., 2014). Abnormalities in OFT-derived structures are the most frequent congenital heart defects (Fahed et al., 2013). In our system, we observed that higher RA dosages promoted aSHF specification towards the RV chamber identity (IRX1, PRDX1, NPPA) (Figures 3A, S3A, and S3B), while the absence of RA signaling promoted the expression of OFT markers (WNT5A, ISL1, HAND2, WNT11, BMP4, RSPO3) but not chamber markers such as NPPA (Figures 3B-C and S3B). We confirmed these observations by immunostaining for ISL1, WNT5A, and HAND2 (Figure 3D, 3D’ and S3C). Consistently, the OFT gene expression profile shows that the marker WNT5A is upregulated at day 4.5 and that OFT cardioids are more mesenchymal-like (Figure S3A), delayed in differentiation (Figure 3B), and are smaller in size compared to RV cardioids (Figure 3E, S3E). Further optimization of WNT pathway inhibitors revealed that C59 led to higher expression of chamber markers (NPPA) in the RV, whereas XAV-939 promoted upregulation of OFT genes (WNT5A, MSX1, BMP4) (Figure S3D). Most OFT cardioid cells are TNNI1+ CMs, and only a few cells show fibroblast or endothelial marker expression (Figure S3F). Thus, after specification, aSHF progenitors are directed into either RV or OFT-like cardioids (OFT cardioids) by the presence or absence of RA, respectively.

**Figure 3:**
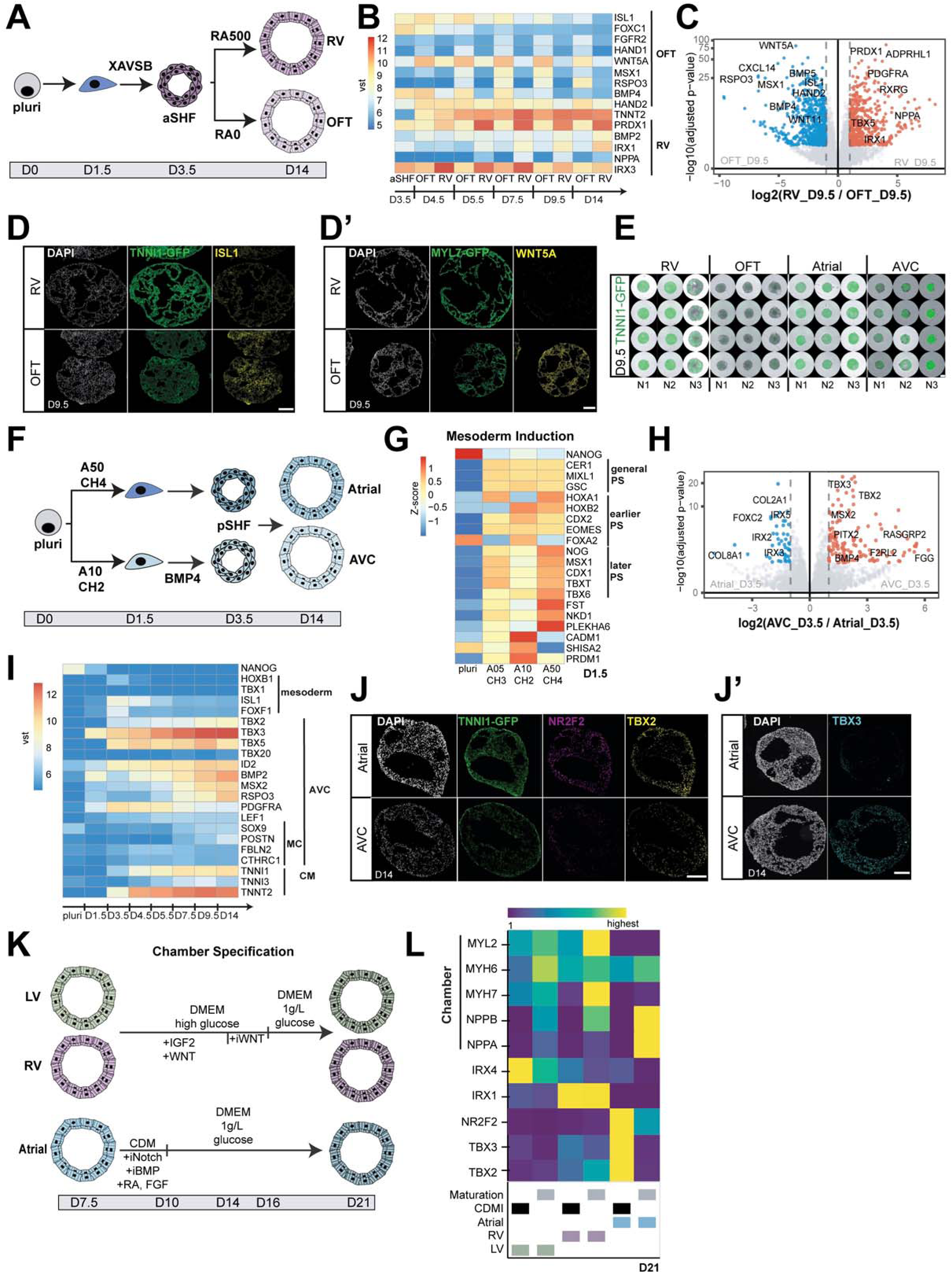
Specification into OFT, AVC, and chamber cardioids. (A) Differentiation protocol of aSHF progenitors into RV and OFT cardioids. (B) Bulk RNA-seq time course of developing RV and OFT cardioids reveal expression of lineage-specific genes. (C) Global gene expression difference of RV and OFT cardioids at day 9.5. (D) Fluorescent cryosection images of RV and OFT cardioids for OFT markers ISL1 (ICC) and (D’) WNT5A (in-situ hybridization chain reaction (HCR)) staining at day 14. (E) Whole-mount images of RV, OFT, atria, and AVC cardioids derived from the TNNI1-GFP reporter line of three biological replicates at day 9.5. scale bar, 500µm (F) Differentiation protocol for atrial and AVC cardioids. (G) Differentially expressed genes by bulk RNA-seq at day 1.5 between AVC, FHF, and SHF progenitors. PS: primitive streak (H) Volcano plot showing most differentially expressed genes by bulk RNA-seq of AVC compared to atrial progenitors at day 3.5. (I) Heatmap showing gene expression by bulk RNA-seq of AVC cardioids over time. MC: mesenchymal (J) ICC images of NR2F2, TBX2, and (J’) TBX3 on cryosections of atrial and AVC cardioids on day 9.5. (K) Chamber specification protocol of LV, RV, and atrial cardioids. (L) RT-qPCR of LV, RV, and atrial cardioids after maturation protocol versus CDM-Insulin on day 21. All scale bars in this figure have a length of 200µm except where specified. All bulk RNA-seq data are the mean of N=3. vst: variance-stabilized transformed counts.

In vivo, pSHF-derived CMs comprise most of the atria and contribute to the AVC, a crucial region in which valves and pacemaker elements develop. In the mouse, pSHF precursors are known to locate in different areas in the primitive streak and will migrate out at different developmental time points (Ivanovitch et al., 2021; Lawson et al., 1991; Tam et al., 1997). pSHF precursors that give rise to the AVC migrate earlier, while the atrial pSHF precursors later. Based on these developmental studies, we hypothesized that mesoderm induction conditions for the two pSHF populations would likely differ in the cardioid platform. Indeed, we found that intermediate Activin and low WNT activation levels during mesoderm induction resulted in higher expression of early primitive streak markers at day 1.5 (Figure 3G), leading subsequently to the upregulation of AVC-specific genes (TBX2, TBX3) and down-regulation of atrial genes (NR2F2) at day 9.5 (Figure S3G and S3H). The pSHF signature (HOXB1, FOXF1, TBX5) at day 3.5 remained in both pSHF populations (Figures 3I and S3H). Another notable difference between AVC and atrial development *in vivo* is the high exposure of the AVC region to BMP ligands. As hypothesized, the addition of BMP4 at the patterning stage upregulated early AVC markers (Figures S3G and S3H). Finally, the combination of the optimized induction and patterning stages (Figure 3F) drove pSHF specification towards AVC identity, as seen by RNA-seq (Figures 3H and 3I), and high protein expression of TBX2 and TBX3 and lower protein expression of NR2F2 compared to atrial cardioids (Figures 3J and 3J’). Moreover, AVC-like cardioids (AVC cardioids) were smaller than atrial cardioids (Figures 3E and S3E), and only a few cells were PECAM1+ or COL1A1+ (Figure S3F). Overall, the sub-specification of pSHF progenitors into atria or AVC cardioids starts as early as the mesoderm induction stage, indicating that pSHF retains developmental plasticity.

During the early stages of cardiogenesis (E8.0 in the mouse), the atria and AVC have similar gene expression profiles (de Soysa et al., 2019), which we also observed in our atrial cardioids that still express the primary myocardium marker TBX2 at day 9.5. *In vivo*, the atria, LV and RV subsequently start upregulating the chamber myocardium gene expression programs, while the AVC and the OFT do not. To achieve chamber specification, we attempted a previously published ventricular CM chamber specification and maturation signaling treatment used in embryoid body differentiations (Funakoshi et al., 2021). Although our data show that LV and RV cardioids upregulated the chamber markers MYL2, MYL7, NPPB, and NPPA (Figure S3I), this approach did not stimulate the atrial chamber program (Figure S3I). In the following steps, we, therefore, tested the effects of different signaling pathway activities to identify the combination of factors that promotes explicitly atrial chamber differentiation. We found that the combination of FGF and RA pathway activation and NOTCH and BMP pathway inhibition promoted the atrial chamber gene program while down- regulating AVC-specific genes (Figure S3J). When we combined this treatment with a low-glucose medium, similar to metabolic maturation treatments (Karbassi et al., 2020), we observed further chamber differentiation (Figures 3K and 3L). Cumulatively, we demonstrate that we can specify and differentiate cardioids into the five major compartment identities found in the embryonic heart.

### Functional characterization of the five cardioid subtypes

The heart must function while developing; thus, understanding early cardiac activity during the formation of the different embryonic heart compartments is imperative to developing experimental multi-compartment heart models. Animal experiments suggest that considerable differences in spontaneous contraction (automaticity) and beating frequency exist between the heart compartments. Still, human embryonic data from 19 to 35 days post-fertilization are entirely missing (Watanabe et al., 2016). The FHF-derived heart tube and early LV region start to contract first (Tyser and Srinivas, 2019; Tyser et al., 2016) but lose automaticity as they mature. In contrast, the developing atria and AVC start to beat later and maintain automaticity until the cardiac pacemaker elements have formed (Christoffels et al., 2010). We hypothesized that the cardioid compartment platform could be used to investigate these early functional developmental differences before the formation of pacemakers and before human *in vivo* data can be acquired.

The cardioid platform is particularly advantageous for investigating contraction dynamics using microscopy because of the ease of imaging cardioids in high throughput. The contraction behavior of day 6.5 compartment-specific cardioids showed automaticity of beating in 90-100% of LV, atria, and AVC, and a great extent of contraction. In contrast, only 18% of the RV cardioids contracted spontaneously, and only 8% of the OFT cardioids contracted with a low contraction extent (Figures 4A, 4C, and S4A). On day 9.5, atria and AVC cardioids maintain automaticity, while automaticity and contraction rate decrease in LV, RV, and OFT cardioids (Figures 4A-C). Similar to *in vivo* development (van Weerd and Christoffels, 2016), the loss of automaticity correlates with the downregulation of the potassium/sodium channel encoding gene HCN4, commonly expressed in pacemakers (Figure 4D). Importantly, these observations were reproducible across both technical and biological replicates (N=2-7, n= 16, resulting in 80, 65, 48, 48, and 33 cardioids, respectively, for LV, RV, OFT, Atria, and AVC) (Figures 4A-C). To gain further insights into signal propagation in cardioids, we generated a GCaMP reporter line to trace calcium transients (Figure S4B). Interestingly, we found that each cardioid subtype has its distinct beating pattern; the atrial and AVC cardioids beat very regularly (100% of beating cardioids), while the LV cardioids beat in regular bursts (95%). The RV cardioids tend to contract once with several sub-contractions in a row followed by relaxation (70%) (Figures S4B).

**Figure 4:**
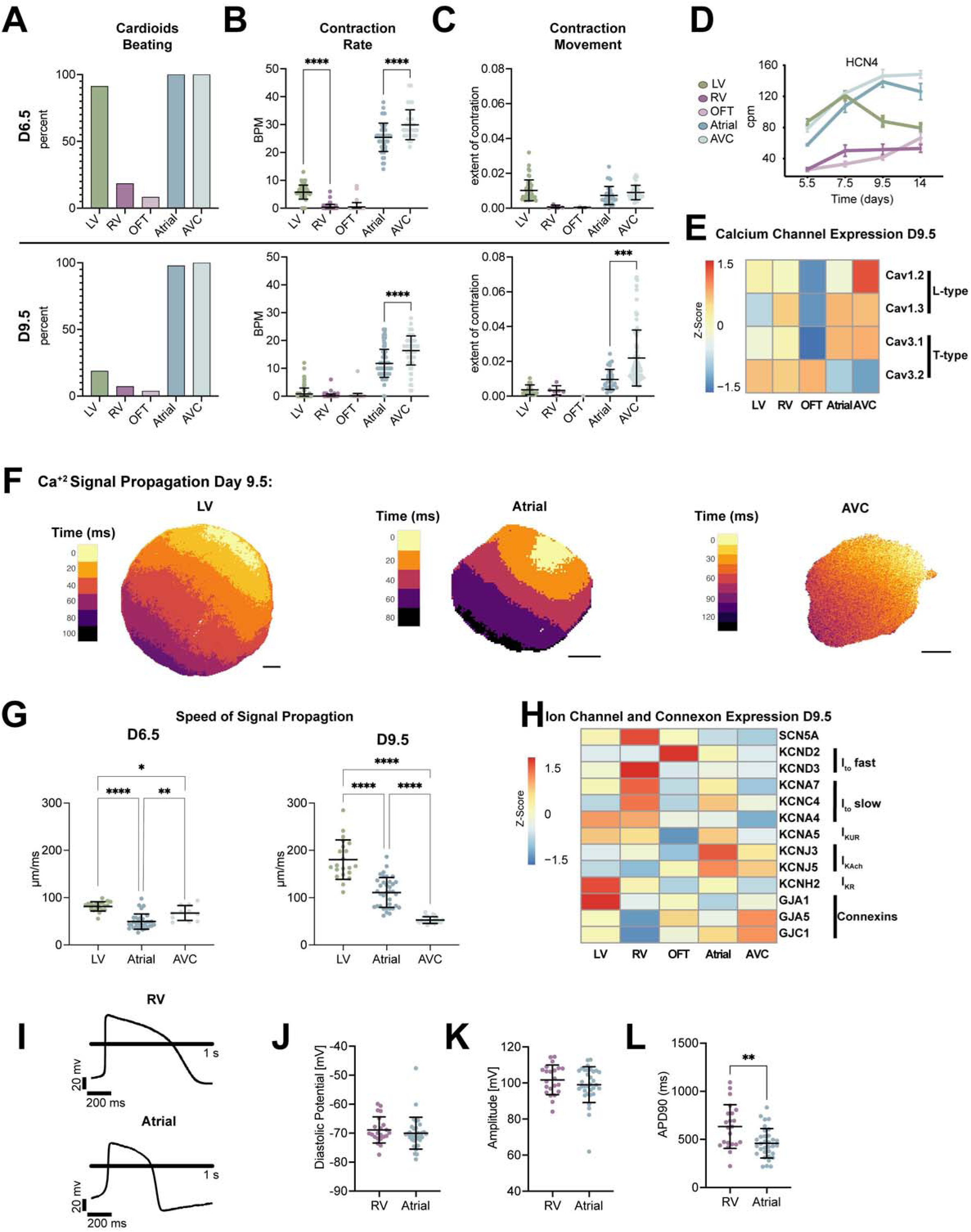
Functional characterization of cardioid subtypes. (A-C): For days 6.5 and 9.5. Experiments were performed in N=2-7, n= 16, resulting in 80, 65, 48, 48, and 33 cardioids for LV, RV, OFT, atria, and AVC, respectively. (A) Quantification of the percentage of cardioids that spontaneously contract within one minute of recording (B) Quantification of beats per min (BPM) of cardioids (C) Quantification of the extent of contraction representing a proxy of how far the cardioid’s edge moves during one contraction. Cardioids that are not beating were excluded. (D) RNA expression by bulk RNA-seq quantification of HCN4 over time. Each dot represents the mean ± SD. cpm: counts per million. (E) Bulk RNA- seq shows the expression of calcium channels of both L and T-types on day 9.5 of differentiation. Legend: Z- score. (F) Representative calcium signal propagation image of LV, atrial, and AVC cardioids for one beat. The color of each pixel corresponds to the time the intensity reached 50% of peak intensity. The black bars underneath are a distance scale bar of 200 µm (G). Quantification of the speed of signal propagation across different cardioid types at day 6.5 (left) and day 9.5 (right). Each point represents the mean speed for all beats of a single cardioid. 21 cardioids from 3 biological replicates for LV, 35 cardioids from 4 biological replicates, and 22 cardioids from 1 biological replicate for AVC were analyzed. (H) Heatmap of bulk RNA-seq at day 9.5 Legend: Z-score. (I-L) Patch-clamp analysis performed on single cardiomyocytes after 9.5 days of differentiation: Each point represents the mean from one cell for 15-20 consecutive APs. 22 cells were analyzed for RV, and 33 cells were analyzed for Atria (I) representative AP (action potential) curves, (J) diastolic potential (K) Amplitude of AP, and (L) Action potential duration (APD90). All graphs show mean ± SD. For all statistics, one-way ANOVA was used except for L, where the student’s T-test was used. *p < 0.05, **p < 0.01, *** p < 0.001, ****p < 0.0001.

We also investigated how calcium transits across the whole cardioid; we found that the LV cardioid has a prolonged transient compared to atrial and AVC cardioids (Figures S4C, S4D, S4E, and S4F) as reported *in vivo* (Koopman et al., 2021). In addition, the speed of signal propagation across cardioids differed between cardioid types and differentiation stages, where the LV cardioid’s calcium transients propagated faster (180.3 um/ms) than both the atrial (111 um/ms) and AVC cardioids (52.48 um/ms) (Figures 4F and 4G). This is consistent with the upregulation of GJA1 (CX43), specifically in LV cardioids which have high conductance, and the upregulation of GJC1 (CX45), in AVC cardioids which have low conductance properties (Figure 4H) as seen in vivo (Christoffels et al., 2010). Within one cardioid, the origin of signal propagation varied between beats (Video S1). We also observed differences between cardioid subtypes, as reflected by expression differences in both T and L-type calcium channels (Figure 4E). Overall, each cardioid subtype has a distinct contraction and signal propagation profile at these early embryonic stages, which are not accessible in humans.

During early heart development, ion channel expression is relatively uniform, but in later stages, chamber-specific gene expression profiles and species-specific action potential (AP) shapes emerge, often characterized by AP duration (APD) (Christoffels and Moorman, 2009). The cardioid subtypes also develop distinct ion channel expressions by day 9.5 (Figure 4H). To characterize how APs of early human CM subtypes differ within cardioids, we used manual patch-clamp and voltage-sensitive dye (FluoVolt) imaging. RV and atrial cardioids were dissociated to perform patch-clamp analysis (Figures 4I). We observed that the AP duration in atrial CMs was shorter than in RV CMs, similar to isolated human primary CMs (Figures 4I and 4L, S4H, and S4I). Furthermore, the diastolic potential was around –70 mV (Figures 4J), closer to *in vivo* CMs than most other iPSC-derived systems (Verkerk et al., 2021). In addition, the upstroke velocity (Figure S4J) and amplitude (Figure 4K) of the action potentials resembled other *in vitro* models. This was confirmed by using a voltage-sensitive dye to measure the behavior of a larger cell population from dissociated cardioids. We found similar trends in APDs where the LV and RV had longer APDs than the atria (Figures S4K-N). Taken together, the electro-chemical signaling of cardioid subtypes is diverse and immature, providing a system to investigate the developmental electrophysiology of early human cardiogenesis.

### Multi-chamber integration of cardioid subtypes

Embryonic cardiac progenitors become functionally specified in neighboring but separate areas. As SHF- derived RV and atrial precursors migrate into the heart tube, they self-sort to remain in separate compartments, a mechanism vital for the heart’s function (Meilhac and Buckingham, 2018). Studying the molecular basis of this sorting mechanism in embryos is challenging. We hypothesized that *in vitro*-derived a/pSHF and FHF progenitors will have the same potential to self-sort as their *in vivo* counterparts. Indeed, when we dissociated developing cardioid subtypes at day 3.5 derived from either H2B-GFP or LMNB1-RFP hPSC reporter lines and then mixed them (Figure 5A), we observed separation within 24 hours (Figures S5B and 5C) that continued on day 7.5 and beyond (Figures 5B and S5A). In contrast, progenitors of the same subtype did not sort upon mixing (Figures 5B’ and 5SA’). The sorting was consistent with the differential Cadherin-expression signature in the different progenitors, reminiscent of *in vivo* findings (Cortes et al., 2018) (Figures 5D, 5D’ and S5C). Labeled progenitors self-sorted one-day post mixing and kept their identity (Figure 5C). They developed into TNNT2+ CMs at day 7.5 (Figure 5B) and retained the appropriate chamber fate (Figure S5D), confirming that the first two stages of differentiation determine lineage identity. These results also imply that co-differentiation of different progenitors was possible from day 3.5 onward. In summary, the multi-chamber cardioid platform can be used to study further the sorting mechanisms that separate the major compartments of the embryonic heart.

**Figure 5:**
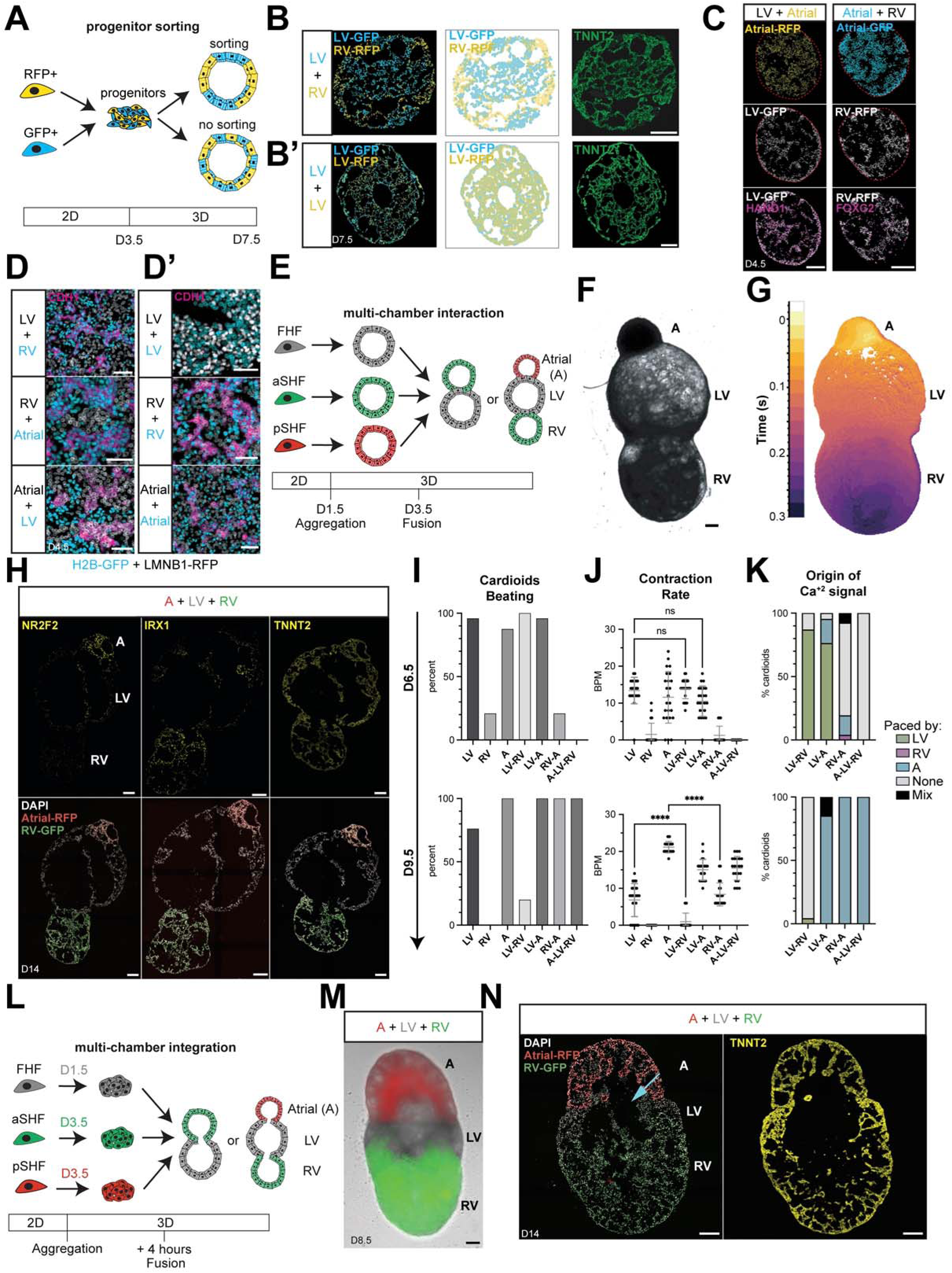
Multi-chamber integration of cardioid subtypes. (A) Schematic of mixing experiments (B-B’) Cross-sections and schematics of cardioids derived from (B) different or(B’) the same cardiac progenitors traced by H2B-GFP or LMNB1-RFP hPSC reporter lines. (C) ICC of (HAND1 (LV) and FOXC2 (RV)) of cardioids one-day post mixing. The red line indicates the edge of the cardioid (D-D’) Zoomed images of differently (D) mixed progenitors and (D’) control in cardioids stained for CDH1 one-day post mixing (day 4.5). (E) Schematic of protocol for multi-chambered cardioids. Different compartments are fluorescently labeled (H2B-GFP and LMNB1-RFP lines). (F) A representative bright-field image from a multi-chamber cardioid on day 6.5 consists of an atrial (A), LV, and RV compartment. (G) Representative calcium signal propagation through a triple chambered cardioid for one beat. The map is colored when each pixel reaches 50% of peak intensity. (H) ICC of cryosection multi-chambered cardioids for NR2F2, IRX1, and TNNT2. Atrial section (A) marked with RFP, LV unmarked, and RV marked with GFP (I) Percentage of cardioids beating for each fusion type, for day 6.5 and day 9.5. (J) Beats per minute (BPM) for the different fusion types on day 6.5 and day 9.5. (K) Origin of calcium signal for multi-chamber cardioid. The color indicates in which cardioid subtype the beat is initiated as a percentage of all cardioids recorded. Mix indicates that different cardioids initiated the beat for each beat, and None indicates that fused cardioids are not beating or that no interaction is happening between the fused cardioids. (I-K) Shown for both day 6.5 and day 9.5. (N=1, n=24) (L) Schematic of protocol for multi-chambered cardioids with shared lumen (M) Representative image of cardioid with three compartments using protocol depicted in L. (O) Representative cross-section of a triple chambered cardioid using the protocol depicted in M stained for TNNT2. Arrow indicates a shared cavity between chambers. All scale bars in this figure have a length of 200µm. All graphs show mean ± SD. For all statistics, one-way ANOVA was used. *p < 0.05, **p < 0.01, *** p < 0.001, ****p < 0.0001. ns: not significant

*In vivo*, the LV, RV, and atrial chambers co-develop seamlessly; however, we lack a multi-chamber model to study this crucial stage and the complex process of cardiac morphogenesis. As cardiac progenitors are specified and sorted already at day 3.5, we hypothesized that co-developing cardioids would also remain separate at this stage but undergo morphogenesis together. When we placed different cardioid subtypes together on day 3.5 (Figure 5E), they co-developed to form a structural connection after 24 hours (Figure S5E). Still, they maintained their distinct identities and compartments (Figures 5F and 5H). In contrast, when we placed cardioid subtypes together on day 5.5, they failed to connect (Figure S5F). Cardioids only co-developed upon fusion on day 3.5, electrochemically connected and contracted in a coordinated manner on day 6.5 (Video S2 and S3; Figures 5G and S5G), demonstrating that the different cardioid subtypes interact functionally. Hereafter, we refer to these structures as multi-chambered cardioids. Multi-chambered cardioids could co-develop in all combinations, allowing us to study the interactions of two-chambered cardioids (Video S2 and Figure S5H) or three-chambered cardioids (atrial, LV, RV fusion) in the same order as within the developing embryonic heart (Video S3; Figures 5F-H and S5G).

Studies in mice and chickens show that directional electrochemical signal propagation occurs early in heart development. Still, this process has not been possible to track in human embryos (Watanabe et al., 2016). Directionality of the electrochemical signal and fluid propagation is established gradually, initially without pacemakers, valves, and septa. The first electrochemical signals are known to appear in the differentiating FHF/LV (Tyser et al., 2016). Once the atrial region develops, it follows the pace of the other areas, ensuring unidirectional signal motion and flow from the atria over the LV to the RV and OFT. To investigate whether the multi-chambered cardioid system recapitulates this process, we measured its calcium signal propagation. We found that each beat originated typically from one location and then propagated through the entire multi-chambered cardioid (Video S3; Figure 5G and S5G), generating unidirectional signal propagation. On day 6.5, most beats originate from the LV region (Figure 5K) and propagate through the RV area, which does not beat independently (Figure 5I). Furthermore, we validated these observations to show that multi-chambered cardioids paced by the LV region maintain the same beat frequency as LV cardioids (Figure 5J). Interestingly, on day 6.5, the pacing potential appears limited as the LV region could not pace the three-chambered cardioids, and the atrial area could not pace a multi-chambered cardioid yet (Figures 5I and 5K). However, as the multi-chambered cardioids developed to day 9.5, the signal originated almost exclusively from the atrial region in all combinations (Figure 5K). Thus, we have demonstrated that multi-chambered cardioids provide a unique tool for deciphering the ontogeny of electrochemical signal propagation throughout the co-developing cardiac chambers.

Heart chambers initially share a lumen before the onset of septation and valve formation (Lin et al., 2012). To explore whether we could model this process *in vitro*, we set out to optimize our protocol further. Strikingly, when we combined early-stage FHF, aSHF, and pSHF cardioids just before the formation of cavities (Figure 5L) (day 1.5 for FHF/LV and day 3.5 for aSHF/RV and pSHF/atrial), we found that they co-developed a shared lumen and still retained cardiomyocyte identity (Figures 5M, 5N, and S5I). Taken together, the characterization of our human compartment-specific and multi-chamber platform supports that it can be leveraged to comprehensively dissect the earliest aspects of human cardiogenesis.

### Mutations cause compartment-specific defects in cardioids

Mutations in genes encoding key cardiac transcription factors (TF) cause compartment-specific congenital defects, where autonomous and non-autonomous effects are difficult to disentangle. Moreover, species-specific variations in TF expression and function are becoming increasingly clear in the embryonic development (Rossant and Tam, 2022). To genetically validate the specificity of the human cardioid compartment platform, we next generated knockout (KO) hPSC lines for the prominent TFs ISL1 (Figure S6A) or TBX5 (Figure S6G) and for FOXF1 (Figure S6J), which is less characterized.

Mutations in ISL1 are known to cause severe cardiac malformations in the OFT and RV, partial defects in the atria, and lethality in mice at E10.5 (Cai et al., 2003). In a time-course analysis of ISL1 KO cardioids from day 2.5 to 9.5, we noted severe impairment of cavity morphogenesis and size at day 5.5 in the OFT and atrial cardioids, while the impact on RV cardioids was more subtle (Figure S6D). On day 9.5, there was a significant size difference between KO and WT in all cardioid subtypes (Figures 6A and S6E). Furthermore, we found that misregulated gene expression was already apparent at day 3.5, as evidenced by lower levels of the TFs MEF2C and MYOCD, indicative of aberrant differentiation progression (Figure S6B)(Gao et al., 2019). We saw the most drastic gene expression changes in the OFT cardioids, with HAND2 and BMP4 being down-regulated (Cai et al., 2003) and TBX5 being up-regulated (Figure S6B). In atrial cardioids, the pSHF marker HOXB1 was downregulated (Figure S6B), while the NR2F2, RSPO3, WNT5A, and MYL7 genes were misregulated across all the different cardioid subtypes at day 9.5 (Figure 6C). The efficiency of CM differentiation was severely affected in the RV KO cardioids, noticeably lower in atrial and OFT KO cardioids, while the LV was less affected (Figure 6B). Although atrial cardioids still maintained their identity, albeit with delayed differentiation and onset of contraction (Figures 6B-D and S6C), major OFT regulators were misregulated, leading to a global gene expression shift from OFT to atrial (NR2F1+/2+, HEY1+, WNT5A-) identity (Figure 6D, 6F, and S6F)(Devalla et al., 2015; Quaranta et al., 2018). Consistent with the gene expression analysis, most ISL1 KO OFT cardioids started beating at a similar rate as atrial cardioids on day 14 (Figure 6E). Thus, the cardioid platform shows the most severe defects for the ISL1 KO in similar compartments as seen *in vivo*, allowing detailed, human-specific, and autonomous dissection of specific effects.

**Figure 6:**
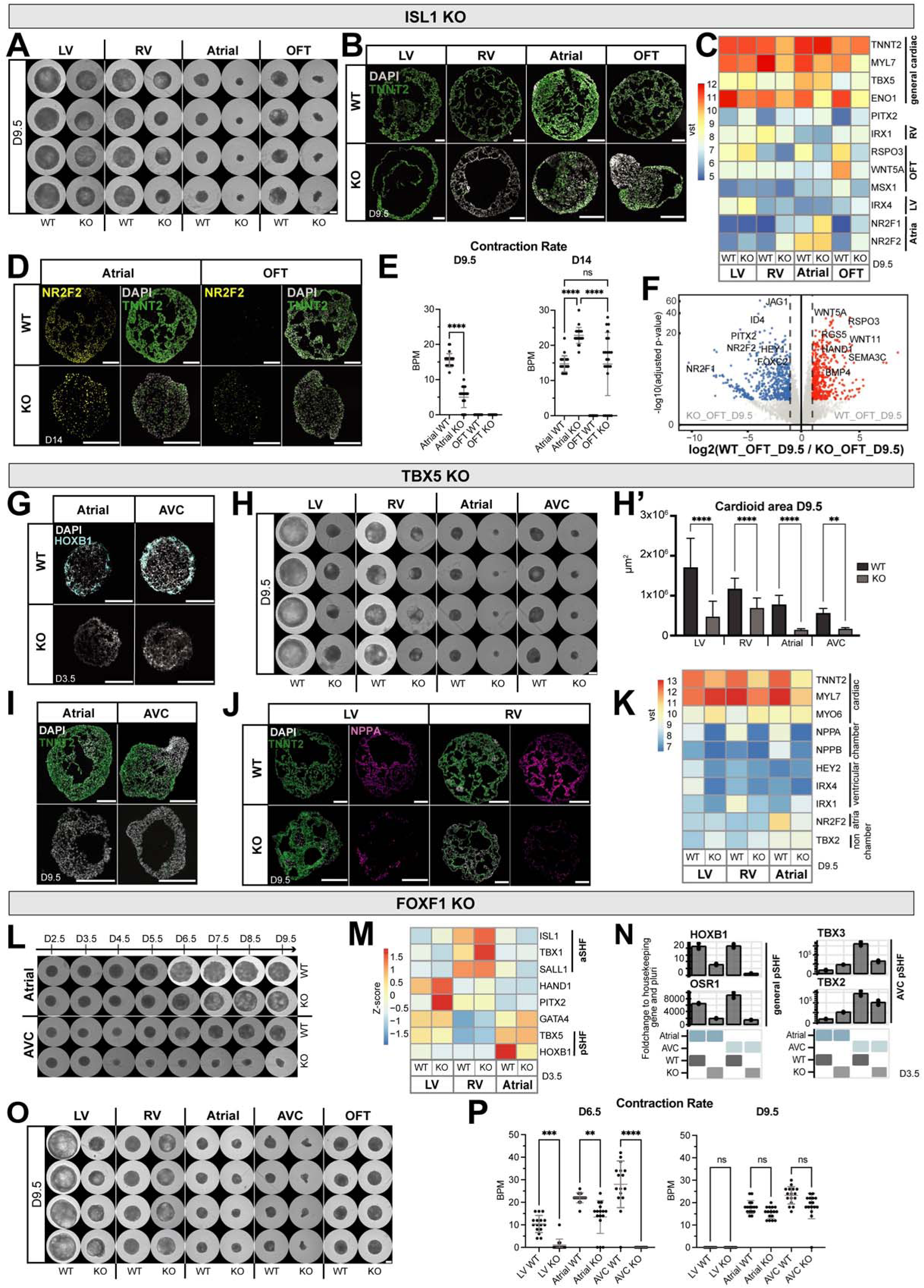
Mutations cause compartment-specific defects in cardioids. (A) Whole-mount images of WT and ISL1 KO cardioids ISL1 KO cardioids using LV, RV, atrial, and OFT protocol at day 9.5. scalebar, 500µm. (B) Immunostaining of TNNT2 on cross-sections of WT and ISL1 KO cardioids at day 9.5. (C) A bulk-RNA-seq analysis shows misregulated genes in ISL1 KO cardioids compared to WT at day 9.5. (D) NR2F2 staining (atrial marker) of atrial and OFT WT vs. ISL1 KO cardioids on day 14. (E) Contraction analysis on days 9.5 and 14 of atrial and OFT WT and ISL1 KO cardioids (N=1, n=24). (F) Volcano plot showing global gene expression differences of OFT cardioids using ISL1 KO compared to WT at day 9.5. (G) RNA-scope staining of HOXB1 (pSHF marker) in atrial and AVC TBX5 KO cardioids compared to WT at day 3.5. (H) Representative whole-mount images of TBX5 KO and WT cardioids and (H’) quantification of the cardioid area at day 9.5 (N=3, n=8). scalebar, 500µm. (I) Immunostaining of CM marker TNNT2 in atrial and AVC TBX5 KO and WT cardioids. (J) Staining of NPPA (chamber marker) and TNNT2 of LV and RV TBX5 KO and WT cardioids at day 9.5. (K) Bulk RNA-seq analysis showing differentially expressed chamber-specific genes (NPPA, NPPB) and identity genes of TBX5 KO compared to WT cardioids at day 9.5. (L) Time course of atrial and AVC cardioids in FOXF1 KO and WT. scalebar, 500µm. (M) Heatmap of bulk RNA-seq dataset of cardiac progenitors at day3.5 showing misregulated genes of FOXF1 KO compared to WT. (N) Representative RT-qPCR at day 3.5 is comparing atrial and AVC FOXF1 KO and WT cardioids. Fold change normalized to a housekeeping gene (PBGD) and pluripotency. (O) Whole-mount images of WT and ISL1 KO cardioids using LV, RV, atrial, and OFT protocol on day 9.5. scalebar, 500µm. (P) Contraction analysis on days 6.5 and 9.5 of LV, atrial, and AVC cardioids, showed reduced BPM at day 6.5 in FOXF1 KO compared to WT (N=1, n=24). vst: variance-stabilized transformed counts. All scale bars in this figure have a length of 200µm. All bar graphs show mean ± SD. For all statistics, one-way ANOVA was used. *p < 0.05, **p < 0.01, *** p < 0.001, ****p < 0.0001. ns: not significant.

TBX5, another prominent cardiac TF, is a critical regulator in FHF and pSHF progenitors and is responsible for driving the chamber gene expression program (Bruneau et al., 2001; Moskowitz et al., 2012). Accordingly, disruption of the TBX5 gene leads to atrial and ventricular septal defects and conduction defects, and when mutated, is associated with congenital heart defects in patients with Holt–Oram syndrome (Bruneau et al., 2001). When we differentiated TBX5 KO cardioids, global gene expression analysis on day 3.5 revealed that a range of aSHF markers (TBX1, FOXC2, FGF10) got upregulated in TBX5 KO atrial cardioids (Figure S6H) and that TBX2/3 were downregulated in AVC cardioids, in line with *in vivo* findings (Figure S6I)(Moskowitz et al., 2012). We also observed that HOXB1 expression was diminished in atrial and AVC KO cardioids at day 3.5 (Figure 6G and S6H). Moreover, TBX5 KOs differentiated into LV cardioids showed an upregulation of HAND2, GATA4, and ISL1 (Figure S6H). In contrast, RV KO cardioids showed no major defects, except for ISL1 upregulation on day 3.5 (Figure S6H). On day 9.5, we observed severe morphogenetic phenotypes in all cardioid subtypes (Figures 6H and 6H’). Atrial and AVC KO cardioids even failed to differentiate into CMs, as seen by the low expression of TNNT2 (Figures 6I and 6K). LV and RV KO cardioids mainly featured inefficient CM differentiation, with downregulation of the chamber-specific marker NPPA (Figures 6J and 6K). All TBX5 KO cardioid subtypes showed a prominent defect in ventricular chamber marker expressions, such as NPPA/B, IRX4/1, HEY2, and upregulation of non-chamber marker TBX2 in RV and LV KO cardioids (Figure 6K), in line with *in vivo* findings (Bruneau et al., 1999). Overall, the TBX5 KO showed specific phenotypes at different stages, while LV, atrial, and AVC KO cardioids were affected already at the progenitor stage and failed to form CMs, LV, and RV KO cardioids featured a mild phenotype at the CM specification stage when chamber-specific genes were downregulated.

Finally, the Forkhead box transcription factor (FOXF1) is a specific regulator of the pSHF lineage (Hoffmann et al., 2014). Disruption of FOXF1 leads mainly to atrial septation defects, but mutant mice already die by E8.0 due to defects in extraembryonic mesoderm, precluding further analysis of cardiac phenotypes. The broad expression of this TF in the early mesoderm is consistent with its proposed role in establishing cardiovascular progenitor identity in the early lateral mesoderm (Kang et al., 2009). When we analyzed FOXF1 KO cardioids in all subtypes, we observed at day 3.5 an earlier onset of cavity morphogenesis in atrial KO cardioids (Figure S6J, yellow arrow). In contrast, the AVC KO cardioids fail to form cavities (Figures 6L and S6L). The main pSHF markers (HOXB1, OSR1) and AVC markers (TBX2, TBX3) were downregulated in FOXF1 KO cardioids (Figures 6M and 6N), consistent with pSHF specification failure. In contrast, only a few genes were misregulated at day 3.5 in LV and RV KO cardioids, including upregulation of PITX2 and TBX1, respectively (Figure 6M). On day 9.5, we observed that the LV and AVC KO cardioids were smaller, while RV, atrial, and OFT KO cardioids were of average size (Figures 6O and S6K). Interestingly, atrial KO cardioids acquired a more ventricular identity (IRX1+, IRX4+, HEY2+, NRF2F2-) and developed more extensive cavities (Figures S6L, S6M, and S6N), while AVC KO cardioids failed to differentiate efficiently (Figure S6L). As expected, we did not observe a severe phenotype in the RV and OFT KO cardioids, except for the downregulation of NPPA in all subtypes (Figures 6O, S6L, and S6M). A less severe phenotype appeared in the LV KO cardioids, where genes involved in cardiac contraction (ENO1) were downregulated (Figure S6M), leading to a lower beating rate, as confirmed by functional assays (Figure 6P). Atrial KO cardioids also showed a lower beating rate, while AVC KO cardioids did not contract at day 6.5 (Figure 6P). These results suggest that FOXF1 has compartment-specific roles, particularly in the pSHF lineage, showing differential effects in atrial versus the AVC cardioids. In summary, the cardioid platform can be employed to dissect human stage- and compartment-specific genetic cardiac defects of specification and morphogenesis without compensatory mechanisms present in the embryo.

### A multi-chamber cardioid platform for screening teratogen-induced heart defects

Beyond genetic causes, congenital heart defects can also be caused by teratogens (e. g. drugs, toxins, metabolites) (Kalisch-Smith et al., 2019). However, no human system exists in which it is possible to investigate, in a high-throughput and easily quantifiable manner, whether teratogens cause compartment- specific defects (Mantziou et al., 2021). Before setting out to test the potential applicability of the cardioid compartment platform in this setting, we first confirmed that a non-teratogenic factor Aspirin (Mantziou et al., 2021; van Meer et al., 2019), at a high 30 μM dosage, did not cause any morphological or significant gene expression differences (Figures S7A, S7A’, and S7B), showing that the cardioid system is not overly sensitive to non-teratogenic drugs. Next, we tested thalidomide, a well-known teratogen in humans, but not rodents (Yamanaka et al., 2021). Because thalidomide binds and interferes with the TBX5 transcriptional activity (Khalil et al., 2017), it causes severe cardiac septal defects, downregulation of chamber markers (NPPA), and limb malformations. When we used the high-throughput cardioid platform to dissect the effects of thalidomide at different concentrations (0.1 μM, 1 μM, and 10 μM compared to human therapeutic plasma concentrations at ∼2.4 μM), we observed dose-dependent impacts in separate compartments. We detected the most striking, low dosage effect on AVC cardioid formation, which was not previously known (Figures 7A and 7A’), Intermediate phenotypes in LV and RV cardioids, and more subtle effects on atrial and OFT cardioids (Figures 7A and 7A’). Gene expression profiles of treated samples revealed a downregulation of the chamber- specific marker NPPA for all lineages except for the RV and OFT and a dosage-specific misregulation of compartment identity markers (NR2F2, IRX1/4) (Figure 7B).

**Figure 7:**
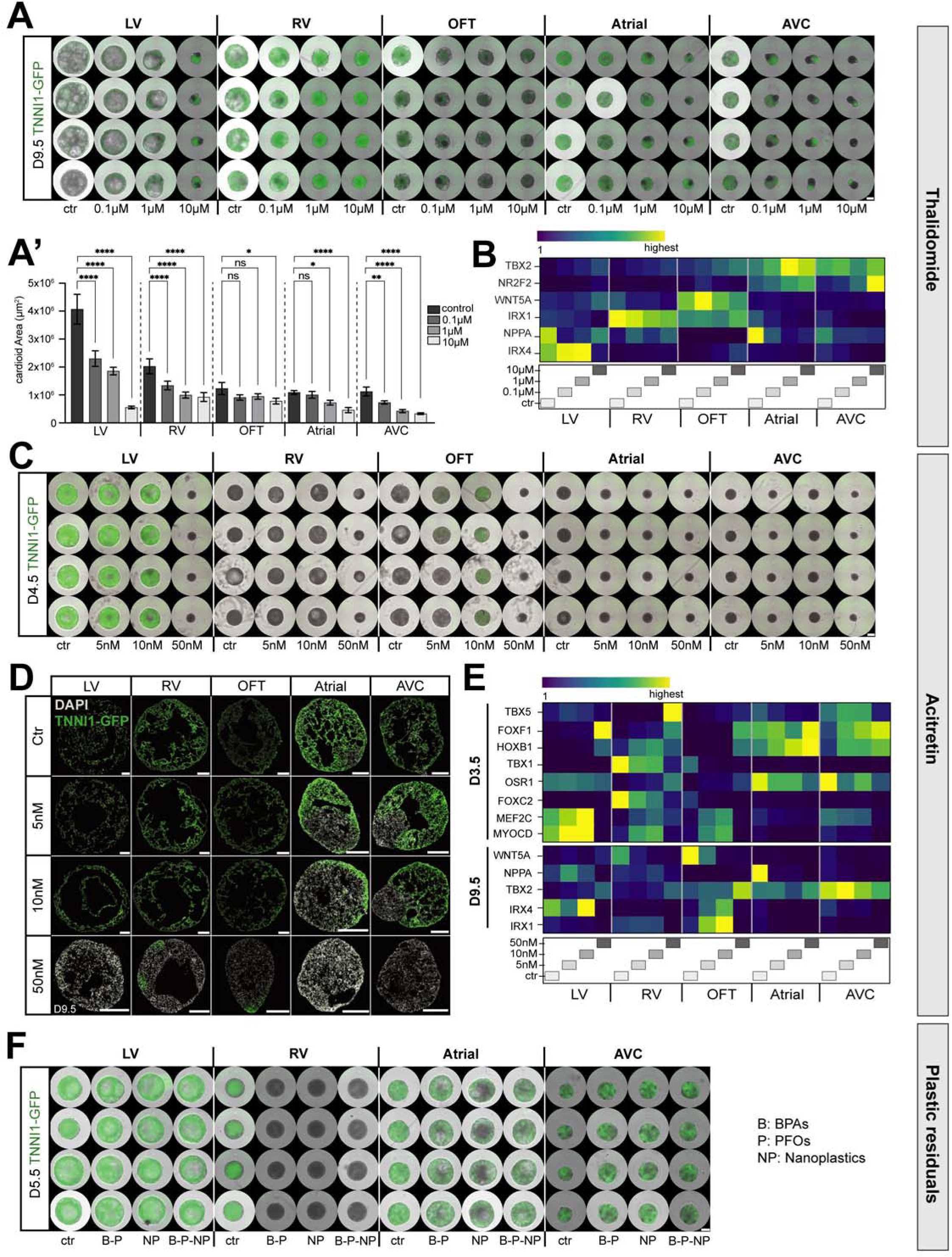
A multi-chamber cardioid platform for screening teratogen-induced cardiac defects. (A) Whole-mount images and (A’) quantification of the size of cardioids derived from TNNI1-GFP reporter line induced with different concentrations of thalidomide compared to control cardioids (day 9.5) (N=1, n=8). ctr: Control. (B) Representative RT- qPCR from day 9.5 cardioids treated with thalidomide showing misregulation of lineage-specific genes. (C) Whole-mount images of cardioids derived from the TNNI1-GFP reporter line induced with different concentrations of acitretin (day 4.5) (N=1, n=8). (D) Cryosection of cardioids derived from TNNI1-GFP reporter line at day 9.5 treated with Acitretin. scalebar, 200µm. (E) Representative RT-qPCR from day 3.5 and day 9.5 cardioids induced with acitretin. (F) Whole-mount images of cardioids derived from the TNNI1-GFP reporter line treated with BPAs (B), PFOs (P), and nanoplastics (NP) compared to control cardioids (day 9.5) (N=1, n=8). All cardioids were induced with teratogens starting from mesoderm induction (day 0) until day 9.5. RT-qPCR: Fold change normalized to a housekeeping gene (PBGD) and pluripotency. All scale bars in this figure have a length of 500µm except where specified. All bar graphs show mean ± SD. For all statistics, one-way ANOVA was used. *p < 0.05, **p < 0.01, *** p < 0.001, ****p < 0.0001. ns: not significant.

Next, we considered retinoid derivatives, used in treatments against leukemia, psoriasis, and acne, as another class of compounds known to induce congenital defects, particularly malformations of the AVC and OFT derivatives (Soprano and Soprano, 1995). Since RA signaling plays a crucial role during heart development and in our cardioid system, we expected that the cardioid compartment platform would allow us to dissect its stage-specific underlying mechanisms. When we tested acitretin and isotretinoin (data not shown), we found that strikingly low dosages (e. G., 5nM acitretin atrial; 1nM isotretinoin in LV) caused severe compartment-specific and stage-specific effects. OFT, atrial, and AVC cardioids had defects in specification, patterning, and morphogenesis when treated with acitretin (Figures 7C, 7D, S7C, S7C’). Surprisingly, when using trans-retinol, we only saw a severe morphological effect on OFT cardioids, while all the other cardioid subtypes were unaffected (Figures S7E and S7E’). In OFT cardioids, retinoids caused downregulation of OFT genes (WNT5A, MSX1, ISL1) and upregulation of ventricular and chamber genes (IRX1, IRX4, NPPA), but not atrial genes (Figures 7E and S7F). Moreover, OFT cardioids treated with retinoids differentiated earlier into CMs (Figure 7C and S7D). These data suggest that the cardioid system is surprisingly sensitive to different retinoid compounds exhibiting drug- and compartment-specific effects.

Finally, we considered that different plastic residues are an emerging and ubiquitous class of compounds in our environment with unknown teratogenic effects (Bojic et al., 2020). In general, *in vivo* effects of plastic residues are very difficult to demonstrate, and until now, a validated system in which to unravel specification and morphogenesis defects in the human heart has not been available. Here, we used our cardioid compartment platform to investigate the effect of different combinations of plastic residues (BPAs, PFOs, and nano plastics). Although LV and AVC cardioids were unaffected by the experimental conditions, we observed a delay in RV cardioid specification and inefficient RV and atrial cardioid differentiation (Figures 7F, S7G and S7H).

Together, these results validate that we can discern, with high sensitivity, early developmental effects of known teratogenic drugs, therapeutic agents, and environmental compounds in a human multi-compartment cardiac platform and relate these to cardiac defects observed in patients. As such, our work has broad implications for studying the effects on human cardiac biology in contexts ranging from therapeutic development to environmental studies.

## DISCUSSION

Recently, several self-organizing human heart models have been reported, including cardiac and cardio- endodermal organoids (Drakhlis et al., 2021; Feng et al., 2022; Lewis-Israeli et al., 2021; Silva et al., 2020). However, because this earlier work did not delineate relationships with aSHF, pSHF, and FHF lineages, the resulting identities, and physiology of the cardiac cell types have remained unclear. As a result, ratios of different CM subtypes and the structures they form *in vitro* have not been controlled and are challenging to relate to the *in vivo* heart. To complement the embryo gold standard model, we developed a signaling-controlled cardioid platform representing all major compartments of the embryonic heart. In this system, the three progenitor populations (aSHF, pSHF, and FHF) give rise to five major cardiac compartments separately or in combination (LV, RV, OFT, atria, and AVC), mimicking selected aspects of early human heart development. The optimized 2D to 3D differentiation approach ensures that homogenous progenitors are specified early, thereby reducing heterogeneity and increasing robustness, which are persistent challenges in the organoid field. We demonstrate that our platform is versatile, highly efficient, reproducible, compatible with multiple cell lines, screenable in high throughput experiments, and comprises single compartment or multi-chamber cardioids.

Several reports describe atrial and ventricular CM differentiated from hPSCs, but whether these originate from the FHF, aSHF, or pSHF lineage has not been determined (Devalla et al., 2015; Lee et al., 2017). *In vivo*, the dosage and timing of signaling are coordinated to drive lineage specification during mesoderm induction in the primitive streak. As mesodermal cells migrate out at different times, they take defined positions within the heart fields. Consistently, we found that specific activation levels of Activin/Nodal and WNT signaling instruct specification into distinct SHF, AVC, and FHF progenitors. Following mesoderm induction, TGF-beta/Nodal signaling inhibition is crucial to determining SHF lineage fate choice, which is consistent with the signaling environment in the anterior region of the embryo (Arkell and Tam, 2012; Nandkishore et al., 2018) but was not highlighted before in a/pSHF specification *in vivo* or *in vitro*. Thus, we hypothesize that TGF-beta/Nodal directly represses a/pSHF genes, thereby promoting FHF specification.

Only highly specific combinations of mesoderm induction and patterning signals allow for mimicking the identities, dynamics, and later functionality of the developing cardiac lineages. For instance, both SHF lineages give rise to differentiated CMs that form cavities with a delay, and the aSHF is more epithelial-like and highly proliferative compared to the FHF (Cortes et al., 2018). In contrast to the aSHF and in agreement with *in vivo* observations, the pSHF encompasses a more diverse range of induction and patterning conditions, resulting in either AVC or atrial phenotypes. This careful mimicking of specification and morphogenesis also led to *in vivo*-like functionality reflected by differences in contractions and dynamics of atrial, AVC, LV, RV, and OFT cardioids, as pacemakers are still absent at this developmental stage. Finally, with its different induction and patterning stages, the platform allows for the dissection and control over progenitor and compartment sorting mechanisms and chamber interactions.

We found that the role of RA signaling in lineage specification was more complex in terms of dosage and timing than expected. Previously, RA was reported to distinguish primarily atrial and ventricular specification in the cardiac mesoderm (Devalla et al., 2015; Lee et al., 2017). Here, we found that the absence of RA signaling is essential for initial aSHF specification and later OFT differentiation. Furthermore, relatively low levels of RA are required for LV specification, and high levels of RA early on for atrial specification. However, the timing is crucial, as the aSHF RV specification requires high levels of RA signaling at later stages.

The teratogenic screening experiments confirmed the critical role of RA signaling dosage, showing the strong effect of retinoids on differentiation speed, specification efficiency and direction, morphogenesis, and physiology, ultimately leading to compartment-specific defects.

Interactions between cardiac lineages during the earliest stages of heart development, including cardiac mesoderm specification, morphogenesis, and functional differentiation, are notoriously difficult to analyze and inaccessible in human embryos. In addition, studies of human embryo development reveal a growing list of differences between species in expression patterns of critical developmental and functionality genes (Cui et al., 2019; Rossant and Tam, 2022; Verheule and Kaese, 2013). Such aspects are key to understanding the human-specific impact of mutations and teratogens on early human heart development and how this causes embryo failure. A significant advance of our work is that the multi-chamber cardioid platform can be used to study how different progenitor populations sort to remain as separate compartments. A second equally important advance is that we can now explore the ontology of contraction signal propagation through the different early stages (days 20-35) of cardiogenesis, another hidden aspect of human heart development. This is particularly important to understand cases of embryonic cardiac failure that have been attributed to faulty specification and morphogenesis, but where defects in early contraction signal propagation between chambers might have been the culprit.

In conclusion, despite decades of experimental and clinical research, the underlying causes of most cardiac defects remain unknown. Potential culprits include still unidentified mutations in regulatory elements such as enhancers, environmental factors such as pollutants, and more complex interactions between genetic and environmental factors, including diet. Previously, we lacked a system to test all these options in a human context with high throughput, encompassing all cardiac compartments. For the first time, the multi-chamber cardioid platform allows us to dissect all these aspects comprehensively and systematically.

## Limitations of the study

Despite its usefulness, the cardioid system has several limitations at this stage of development. This work focuses on comprehensive modeling of early specification, morphogenesis, and signal contraction propagation of the human embryonic heart. However, we have not modeled processes such as a/pSHF progenitor migration and heart looping, nor interaction with the endoderm where other complementary *in vitro* systems (Drakhlis et al., 2021; Rossi et al., 2019; Silva et al., 2020) might be more suitable to compare to the embryo. Moreover, later stages and processes during heart development have not been represented yet in cardioids, including forming valves, septation, pacemakers, chamber trabeculation and ballooning, coronary vasculature and circulation, and the general growth of the heart. Some congenital defects and many adult-onset cardiomyopathies would require models including these features. Therefore, the multi-chamber platform has been validated mainly using mutations and teratogens affecting the earliest stages while providing at the same time a solid basis for further developments.

## Supporting information

Cardioid showing different origins of signal propagation

3-chambered cardioid showing the direction of signal propagation

Two-chambered cardioid contracting in a coordinated manner

## ACKNOWLEDGEMENTS

We thank all laboratory members for their help, discussions and Katarzyna Warczok for lab management. We are grateful to the VBC Histology, EM and NGS, IMP/IMBA Core, and IMBA SCCF facilities for their services; the Allen Institute for cell lines; We are grateful to Daniel Gerlich for comments on the manuscript, and Life Science Editors for scientific editing. This work was funded by the Austrian Academy of Sciences (OEAW) and the Research Promotion Agency (FFG).

## DECLARATION OF INTEREST

The Institute for Molecular Biotechnology (IMBA) filed a patent application on multi-chamber cardioids with C.S., A.D. T.I., and S.M. named as inventors. P.H. and S.M. are co-founders of HeartBeat.bio AG, an IMBA spin-off company, that aims to develop a cardioid drug discovery platform.

## INCLUSIVITY AND DIVERSITY STATEMENT

We worked to ensure diversity in experimental samples by selecting the cell lines. While citing references scientifically relevant for this work, we also actively worked to promote gender balance in our reference list.

## METHODS

### Cell lines

The WiCell Institute (USA) provided human H9 (female) ES cell lines. The WTC iPS cell line (male, skin fibroblast-derived) was developed at Dr. Bruce R. Conklin’s laboratory (Gladstone Institute of Cardiovascular Disease, UCSF, USA) and purchased from the Coriell Institute for Medical Research (USA). The Allen Institute for Cell Science’s reporter cell lines is derived from the WTC cell line and received from the Coriell Institute for Medical Research (USA).

The E8 culture system (Chen et al., 2011) was used to cultivate all human pluripotent stem cell lines in a customized in-house medium. 0.5 percent BSA (Europa Biosciences, #EQBAH70), in-house manufactured FGF2, and 1.8 ng/ml TGFb1 were added to the original E8 mix (R&D RD-240-B-010). Cells were cultured on Vitronectin XF (Stem Cell Technologies, #7180) coated Eppendorf (Eppendorf SE, #0030 721.110) or TPP (TPP Techno Plastic Products AG, #92012) tissue culture-treated plates and passaged every 2-4 days at approximately 70 percent confluency using Try-pLE Express Enzyme (GIBCO, #12605010). The absence of Mycoplasma contamination in cells was regularly tested.

### Generation of ISL1, TBX5, and FOXF1 knock-out cell lines

ISL1, TBX5, and FOXF1 were knocked out in WTC cells using CRISPR/Cas9 multi-guide sgRNAs (Synthego) for target sites on Exon 3 for ISL1, Exon 5 for TBX5, and Exon 1 for FOXF1 (Figures M1I-K). Cells were transfected using the P3 Primary Cell 4D-Nucleofector X Kit S (Lonza-BioResearch, #: V4XP-3032) and Amaxa 4D- Nucleofector (Lonza-BioResearch). Post nucleofection, cells were incubated in E8 supplemented with 5μM Y- 27632 (Tocris, #72302) on a 6-well plate previously coated with Vitronectin XF (StemCell Technologies, #7180). After two days, the medium was changed to E8 without Y-27632 every other day.

Once cells were approx. 70% confluent, single-cell seeding was performed, and the rest of the cells were collected for gDNA extraction. Successful editing was first assessed on a pool level using agarose gels and Sanger sequencing. Subsequently, single colonies were picked and genotyped to confirm a knockout. Colonies were collected with the help of a microscope (EVOS) and transferred into a pre-coated 96-well plate (Corning, Cat #CLS3370) with 150µl E8/well supplemented with 5µM Y-27632 and Antibiotic-Antimycotic. Genome editing on a pool and clonal level was analyzed using Synthego’s online tool ICE (https://ice.synthego.com/<u>#/</u>).

### Cardioid generation

hPSCs (WTC or H9 lines) are seeded in a 24-well plate (TPP, #92024) at 30-40k cells per well in E8 + ROCKi (5 µM Y-27632, Tocris #1254). All differentiation media are based on CDM that consists of 5 mg/ml bovine serum albumin (Europa Biosciences, #EQBAH70) in 50% IMDM (Gibco, #21980065) plus 50% F12 NUT-MIX (Gibco, #31765068), supplemented with 1% concentrated Lipids (Gibco, #11905031), 0.004% monothioglycerol (Sigma, #M6145-100ML) and 15 µg/ml of transferrin (Roche, #10652202001). 24 hours after seeding in the 24-well plate, the cells are induced with mesoderm induction media. Mesoderm induction media is made up of CDM containing FGF2 (30 ng/ml, Cambridge University), LY294002 (5 µM, Tocris, #1130), Activin A (specific concentrations for different cardioid subtype, Cambridge University), BMP4 (10 ng/ml, R&D Systems RD-314-BP-050), and CHIR99021 (specific concentrations for different cardioid subtypes, see below, R&D Systems RD-4423/50). After 36-40 hours, cells are dissociated with TrypleE (Gibco, #12605010) and seeded in a Corning ultra-low attachment 96 well plate (Corning, #7007) at 15-20k cells/ well in *Cardiac Mesoderm Patterning Media One* made up of CDM containing ROCKi and for all protocols besides the LV cardioids 1 µg/ml of insulin (Roche, #11376497001) plus specific factors depending on cardioid subtype (see below). After seeding, the cells are spun down in a centrifuge for 4 mins at 200g. This protocol is termed 2D-3D standard protocol, used in all Figures unless otherwise specified. Alternatively, hPSCs were seeded into Corning ultra-low attachment 96 well plate with a density of 5000 cells/well. Cells were seeded in a volume of 200 ml containing E8 + ROCKi and collected by centrifugation for 5 minutes at 200 g (Figures 1B, 1C, S1A, and S1B). As another option, 2500 cells/well were seeded directly into induction media +ROCKi (Figure S1F) and collected by centrifugation for 5 minutes at 200 g. For both protocols, cells were induced with mesoderm induction media as described for the 2D->3D protocol. These were termed 3D protocols. For both protocols, 2D-3D and 3D, at day 2.5, the cells are fed with *Cardiac Mesoderm Patterning Media One*. For the next two days, the medium is changed to *Cardiac Mesoderm Patterning Media Two,* made up of CDM containing specific factors depending on the cardioid subtype (see below) and exchanged daily. For the subsequent two days, media is exchanged every day with *Cardiomyocyte Differentiation Media* CDM medium containing BMP4 (10ng/ml), FGF2 (8 ng/ml), and insulin (10 µg/ml)). This medium was termed *Cardiomyocyte Specification Media* (Hofbauer et al., 2021b). For the subsequent days of culture, media is exchanged every other day with CDM containing insulin (10 µg/ml).

Alternatively, the whole protocol can be done in 2D completely by seeding 80,000 – 170.000 cells/24well coated with vitronectin and adding the medium on the same timeline as the cardioids (Figure 1B, S1F). This was termed 2D differentiation.

### Specific media composition of cardioid subtypes Chamber specification protocol

These two sections will be updated soon, and a separate protocol paper will in detail describe the procedure.

### 2D Endothelial cell differentiation

hPSCs were seeded at 100,000 cells/24well coated with vitronectin in E8 medium with 5 mM ROCK-i added. The following day, cells were induced with FLyAB and 1-3 mM CHIR99021 for H9 cells and incubated for 36 – 40 hours. For the next two days, the medium was exchanged to their respective *Cardiac Mesoderm patterning media 1* for the FHF, aSHF, and pSHF. After that, CDM with 200 ng/ml VEGF (200 ng/ml, Peprotech, #AF-100-20) and 2 mM Forskolin (Sigma-Aldrich, #F3917) was given for 2 days, and then the cells were cultured for 1 day in CDM with 100 ng/ml VEGF.

### Mixing of progenitors

Cardiac differentiation of different progenitor cell populations (FHF, aSHF, and pSHF) was done in 24 well plates coated with vitronectin until day 3.5 (2D differentiation). Cell populations were labeled using different colored cell lines (WTC: H2B-GFP, WTC: LMNB1-RFP). On day d3.5, progenitor cells were dissociated by adding 200 ul Try-pLE Express Enzyme (GIBCO, #12605010) for 3 - 4 min at room temperature. Dissociation was stopped by adding 1 ml of CDM containing ROCKi (5 mM). After centrifugation for 4 min at 130 g, cells were resuspended in CDM containing ROCKi (5 mM). Then, two progenitor populations were mixed by seeding 15000 - 20000 cells per progenitor population into ultra-low attachment (corning) into *Co-development Patterning Media*.

### Generation of multi-chambered cardioids

For fusion of two chambers, developing cardioids were transferred on day 3.5 using wide opening tips from individual wells of the 96-well Corning ultra-low attachment plate to sharing wells with one other desired cardioid subtype. This can be accomplished with any combination of LV, RV, or atrial cardioids. For this type of fusion, cardioids were put together in the *Co-development Patterning Media* (Figures 6E-K, 6SE-H; Videos S2, S3). Alternatively, LV progenitors on day 1.5 in 2D could also be combined with RV or atrial progenitors on 2D of day 3.5 in *Co-development Patterning Media* to get a multi-chambered cardioid with at least one shared cavity (Figure 6L-N and S6I). *The* two-chamber/multi-chamber cardioids co-develop if fused at these early stages. Later fusion (e. g. from day 5.5 on) will impair the formation of a shared cavity (Figure S6F).

For the fusion of three cardioids, molds were created with a shape to place the early cardioids that are to be fused in contact with each other in the order as in the natural heart (e.g., a linear order). On day 3.5 of cardiac differentiation, the cardioids were transferred to the molds in a 10cm dish filled with *Co-development Patterning Media* using wide opening tips. Using molds, the cardioids could be arranged in the desired orientation (e. g., first atrial, then LV and RV cardioids, as *in vivo*). Media was not changed while cardioids were fusing in the molds from day 3.5-5.5. On day 5.5, the fused cardioids were moved back to the 96-well plate and media change continued as described above.

To track which cardioids in the fusions, arise from which cell population colored cell lines (WTC: H2B-GFP, WTC: LMNB1-RFP) or dyes were used. For this, cells were stained for one hour before induction using SP- DiIC18(3) (Invitrogen, #D7777) to fluoresce at 564nm or DiIC18(5) (Invitrogen, #D12730) to fluoresce at 668nm.

### Molds for multi-chamber cardioids

Embedding molds have been designed in Tinkercad and were adjusted in diameter and length based on the cardioid size on the day of fusion. Files were exported as .stl files and loaded into the slicer software XYZ print 1.4.0. The negative was printed using transparent PLA with 100% infill density and 0.1mm layer height, and 215°C nozzle temperature. After printing, the negative was treated with a Heatgun (Bosch Hot Air Blower 1800W) at 550°C to carefully melt the surface of the negative, create a smooth finish and remove the 3D printing typical rough surface (Figure M1A-H).

The positive was then cast using polydimethylsiloxane (PDMS). In brief, 5ml of curing agent and 45ml of Monomer (both Sylgard® 184 Elastomer Kit, VWR) were mixed intensively. The mixture was then spun down to remove air bubbles and directly used.

To reduce the extent of bubbles formed during curing, the molds were cast at a low temperature (40°C). For this, the negative was placed into a 10cm dish and slowly covered with 30ml of the liquid PDMS mixture. The negative was then carefully removed from the polymerized PDMS, and residual PDMS was cut off using a scalpel. The mold was then stuck to the bottom of a clean 10cm dish using about 5ml of PDMS and cured at 40°C. To sterilize the mold, it was washed in 70% Ethanol for about 30min in the fume hood with UV turned on. For positioning cardioids in the mold, the mold was rinsed once with PBS and then coated with an anti-adherence rinsing solution (StemCell Technologies, # 07010) to increase the non-stick behavior of the PDMS further. After coating, the molds were rinsed once with PBS and were then ready to use.

### Cryosectioning

Cardioids were fixed with 4% PFA in PBS and cryoprotected with 30% sucrose in PBS before embedding. The embedding was carried out using the O.C.T. cryo embedding medium (Scigen, #4586K1). Embedded tissues were frozen using a metal surface submerged in liquid nitrogen and stored in a −80°C freezer until sectioning on a Leica cryostat. Sections were collected on SuperFrost Plus slides (Thermo Fisher Scientific, #10149870) and kept at −20°C or −80°C until immunostaining.

### Immunostaining

To remove O.C.T., fixed specimens were washed in 1X PBS for 15 min. Optionally tissues were placed in permeabilization solution 0.5% Triton-X100 (Sigma-Aldrich, #T8787) for 5 mins to increase antibody permeabilization. Tissues were then incubated in blocking solution (PBS (GIBCO, #14190094) with 4% donkey serum (Bio-Rad Laboratories, #C06SB) and 0.2% TritonX-100 for at least 30 min. Subsequently, specimens were incubated for 3 hours at room temperature or overnight at 4°C in a blocking solution containing the primary antibody. Then, a 20 min washing in PBS with 0.1% Tween20 (Sigma-Aldrich, #P1379) was performed, followed by incubation for 1 hour at room temperature in a blocking solution containing the secondary antibody. Finally, tissues were washed in PBS with 0.1% Tween20. Slides were mounted using a fluorescence mounting medium (Dako Agilent Pathology Solutions, #S3023) and covered with a cover slip (Menzel-Gläser, #631-0853 VWR).

### RNAscope and In Situ Hybridization Chain reaction (HCR)

RNA-scope was performed with the ACDBio (https://acdbio.com) Manual assay kit using RNAscope Probe-hs-TBX1-C2 (Target region: 100 - 769) and RNAscope Probe-hs-HOXB1-C2 (Target region: 528 - 2015) according to the manufacturer’s instructions. RNAscope Probe-hs-PPIB-C1 was used as a positive control. The probes were designed and manufactured by ACDBio.

HCR fluorescent in situ was carried out using the HCR kit (v.3), purchased from Molecular Instruments (molecularinstruments.org), according to the manufacturer’s instructions with the slight modification of adding 100 μg/ml salmon sperm DNA to the pre-amplification solution and the amplification solution including the hairpins to reduce nonspecific binding. The HCR probe WNT5A (B3) was designed and manufactured by Molecular Instruments.

### Image acquisition and analysis

Spinning disk confocal microscopes (Olympus spinning disk system based on an IX3 Series (IX83) inverted microscope, equipped with a Yokogawa W1 spinning disc) were used to image fixed tissue sections. Live imaging was carried out using an inverted widefield microscope for brightfield and fluorescence (Axioobserver Z1 equipped with an sCMOS camera (Hamamatsu Orca Flash 4). Cardioids in 96-well plates were also imaged using a Celigo Imaging Cytometer microscope (Nexcelom Biosciences, LLC).

All images were analyzed with custom-made scripts created for the Fiji software (Schindelin et al., 2012) and MATLAB (MathWorks).

### Flow cytometry

Cardioids (8 cardioids per condition) were dissociated using a 1.5 mL CM dissociation medium (Stem Cell Technologies, #05025) for 7 - 10 min at 37°C. Dissociation of CMs was stopped by adding 7.5 ml of the support medium. After centrifugation for 4 min at 130 g, cells were resuspended in 600 µl PBS with 0.5 mM EDTA (Biological Industries, #01-862-1B) and 10% FBS (PAA Laboratories, #A15-108). Cells were acquired with a FACS LSR Fortessa II (BD) and analyzed with FlowJo V10 (FlowJo, LLC) software. FACS sorting was performed using a Sony SH800 Cell Sorter (Sony Biotechnology).

### RNA extraction and bulk RNA-seq preparation and analysis

RNA was isolated using an in-house RNA bead isolation kit semi-automated using KingFisher devices (KingFisher Duo Prime). Using the QuantSeq 30 mRNA-Seq Library Prep Kit FWD (Lexogen GmbH, #015), the bulk RNA-seq libraries (N=3, n=8) were generated according to the manufacturer’s instructions. After the preparation of the libraries, samples were checked for an adequate size distribution with a fragment analyzer (Advanced Analytical Technologies, Inc). Then the RNA-seq library was submitted to the Vienna Biocenter Core Facilities (VBCF) Next-Generation-Sequencing (NGS) facility for sequencing. Reads were preprocessed using umi2index (Lexogen) to add the UMI sequence to the read identifier, and trimmed using BBDuk v38.06 (ref = polyA.fa.gz,truseq.fa.gz k = 13 ktrim = r useshortkmers = t mink = 5 qtrim = r trimq = 10 min length = 20). Reads mapping to abundant sequences included in the iGenomes NCBI GRCh38 references were removed using bowtie2 v2.3.4.1 alignment. The remaining reads were analyzed using genome and gene annotation for the GRCh38 assembly obtained from Homo sapiens Ensembl release 94. Reads were aligned to the genome using star v2.6.0c, and reads in genes were counted with featureCounts (subread v1.6.2) using strand-specific read counting (-s 1). Differential gene expression analysis on raw counts, and principal component analysis on variance stabilized, transformed count data were performed using DESeq2 v1.18.1.

### Real-Time Quantitative polymerase chain reaction

The isolated RNA was reverse transcribed to cDNA using the Reverse Transcription Kit (Invitrogen, #18080044) with a C100 Touch Bio-Rad Thermal Cycler. Quantitative PCR was performed using the GoTaq qPCR master mix 2x (Promega, #A6001) with a Bio-Rad CFX384 Real-Time thermal cycler. Values of gene expression of each sample were obtained in triplicates. The Log-fold change of the sample from PBGD as a housekeeping gene and a pluripotent stem cell sample for normalization was calculated using a custom-made script written in python. Primer pairs are specified in Table S1. The most significant fold change for each gene in the heatmaps is specified in Table S2.

### Contraction Analysis

Cardioids were fed fresh CDMI media 1-2 hours before recordings. The 96-well plate was placed in an environmentally controlled stage incubator (37◦C, 5% CO2, water-saturated air atmosphere, Okolab Inc, Burlingame, CA, USA). Each well was imaged using widefield phase-contrast microscopy (Axioobserver Z1 (inverted) with sCMOS camera, Zeis) at 100 frames per second for 30-60 seconds. Videos were then analyzed using MUSCLEMOTION; the data was read into a custom-made software for reported calculations. Percent beating was defined by if the cardioid beat once within the entirety of the recording. Beats per minute were calculated by counting the total number of beats in the video, dividing them by the length of the video in seconds, and multiplying by 60. The extent of contraction is the amplitude given from MUSCLEMOTION divided by the size of the cardioid.

### Calcium Transients

To generate a WTC line expressing the GCaMP6f gene, an AAVS1-integrating construct with a CAG promoter followed by the GCaMP6f sequence was chosen (Mandegar et al., 2016) and introduced as previously described (Hofbauer et al., 2021b).

Cardioids were differentiated into LV, RV, atrial, and AVC or multi-chamber cardioids using the protocol above. Cardioids were fed fresh CDM-I media 1-2 hours before recordings. The 96-well plate was placed in an environmentally controlled stage incubator (37°C, 5% CO2, water-saturated air atmosphere, Okolab Inc, Burlingame, CA, USA). Each well was imaged using widefield microscopy (Axioobserver Z1 (inverted) with sCMOS camera, Zeis) at 50-100 (optimally 50) frames per second for 30-60 seconds. Cardioids were excited at 470 ± 10 nm using a light-emitting diode (LED). Videos were then analyzed using custom-made software. Analysis of signal propagation: The peaks were identified using the whole cardioid analysis pipeline. Only pixels with a maximum intensity higher than an organoid-specific threshold were considered. The intensity was calculated per pixel, normalized to 1, and each trace smoothed using a rolling average over 3 frames. The frame at which a pixel reached 50% of peak intensity was recorded. The first frame in which more than 30 pixels reach 50% max intensity is defined as the first frame. The last frame in which all besides at most 30 pixels reach 50% max intensity is defined as the last frame. The average position of the biggest connected component of pixels which reached 50% of peak intensity first, is considered the origin of signal propagation. The speed of signal propagation is then calculated for all of the other pixels by dividing the distance between the pixel and the origin and the frame difference between the frame where the pixel reaches 50% of peak intensity and the origin frame. The speeds are all averaged together for every pixel and across all beats to determine the speed of signal propagation in the organoid. Images of signal propagation are made using the same technique, and each pixel is color-coded based on frame difference. Cardioids were excluded from this analysis for 4 reasons (1) the cardioid does not beat, (2) there is no clear directionality, (3) there are 2 origins (4) if less than 10% of the cardioid is expressing reporter protein. A list of which cardioids are excluded is listed in supplement Table S3.

### Patch-clamp recordings of single cardiomyocytes

Cardioids were dissociated using the STEMdiff Cardiomyocyte Dissociation Kit (Stem Cell Technologies, #05025) according to the manufacturer’s protocol (incubated for 10 – 20 minutes at 37°C to thoroughly dissociate organoids) and subsequently seeded at low densities of 15 – 40k cells in Laminin-511 E8 Fragment (AMSBIO, #AMS.892 011, 0.5 µg/cm²) coated 35 mm tissue culture-treated dishes (Corning, #430165).

Cells were maintained at 37°C in a humidified incubator with 5% CO2, and whole-cell patch-clamp experiments were performed on single beating cardiomyocytes 4 – 13 days post-plating. Glass micropipettes with resistances of 1.5 – 4 MΩ were pulled from glass capillaries (Harvard Apparatus, #BS4 64-0792) using a Sutter P-1000 Micropipette Puller (Sutter Instrument). The extracellular solution consisted of the following (in mM): 140 NaCl, 5.4 KCl, 2 CaCl2, 1 MgCl2, 5 glucose, and 10 HEPES, with pH adjusted to 7.4 using NaOH. The intracellular pipette solution contained the following (in mM): 150 KCl, 5 NaCl, 2 CaCl2, 5 EGTA, 10 HEPES, and 5 MgATP, with pH adjusted to 7.2 using KOH. Data was acquired at 10 kHz and low pass filtered at 2.9 kHz using a HEKA EPC 10 USB Quadro (HEKA Elektronik GmbH) employing PATCHMASTER NEXT software (HEKA Elektronik GmbH). Spontaneous electrical activity was recorded in current-clamp mode and analyzed using custom-made MATLAB (MathWorks) software. Action potential amplitudes were measured from peak to maximum diastolic potential, and APD values were calculated from action potential peak to the respective percentage of the amplitude’s repolarization. Parameters were individually calculated for 15 – 20 consecutive action potentials per cell and then averaged.

### Optical action potentials

Cardioids were dissociated the same way for patch-clamp experiments, see the previous section, and seeded at 40k cells per well into 96 well-plate (Greiner Bio-One, #655182) wells previously coated with Laminin-511 E8 Fragment (AMSBIO, #AMS.892 011, 0.5 µg/cm²). Cells were kept at 37°C in a humidified incubator with 5% CO^2^ for 7 to 11 days, and the medium was exchanged every two to three days.

CM monolayers were loaded with 0.7 times the manufacturer’s suggested amount of the voltage-sensitive dye Fluovolt (FluoVolt™ Membrane Potential Kit (Thermo Fisher Scientific, #F10488)) after three repeated wash steps with Hank’s Balanced Salt Solution (HBSS, Gibco, #14175-053). Loading was performed at room temperature for 30 minutes, after which the cells were washed with HBSS three more times. The 96-well plate was then placed in an environmentally controlled stage incubator (37°C, water-saturated air atmosphere, Okolab Inc, Burlingame, CA, USA), and fluorescence signals were recorded at an excitation wavelength of 470 ± 10 nm using a light-emitting diode (LED), and emitted light was collected by a photomultiplier (PMT, Cairn Research Ltd. Kent, UK). Fluorescence signals were digitized at 10 kHz. 20 s recordings were subsequently analyzed offline using a custom-made MATLAB (MathWorks) software. APDs were measured at 30%, 50% and 90% repolarization. APD values were calculated from the action potential peak to the respective percentage of the amplitude’s repolarization. Parameters were individually calculated for all recorded action potentials per well and then averaged. The number of analyzed action potentials per well typically ranged between 5 and 20.

## SUPPLEMENTARY FIGURES LEGEND

**Figure S1:**
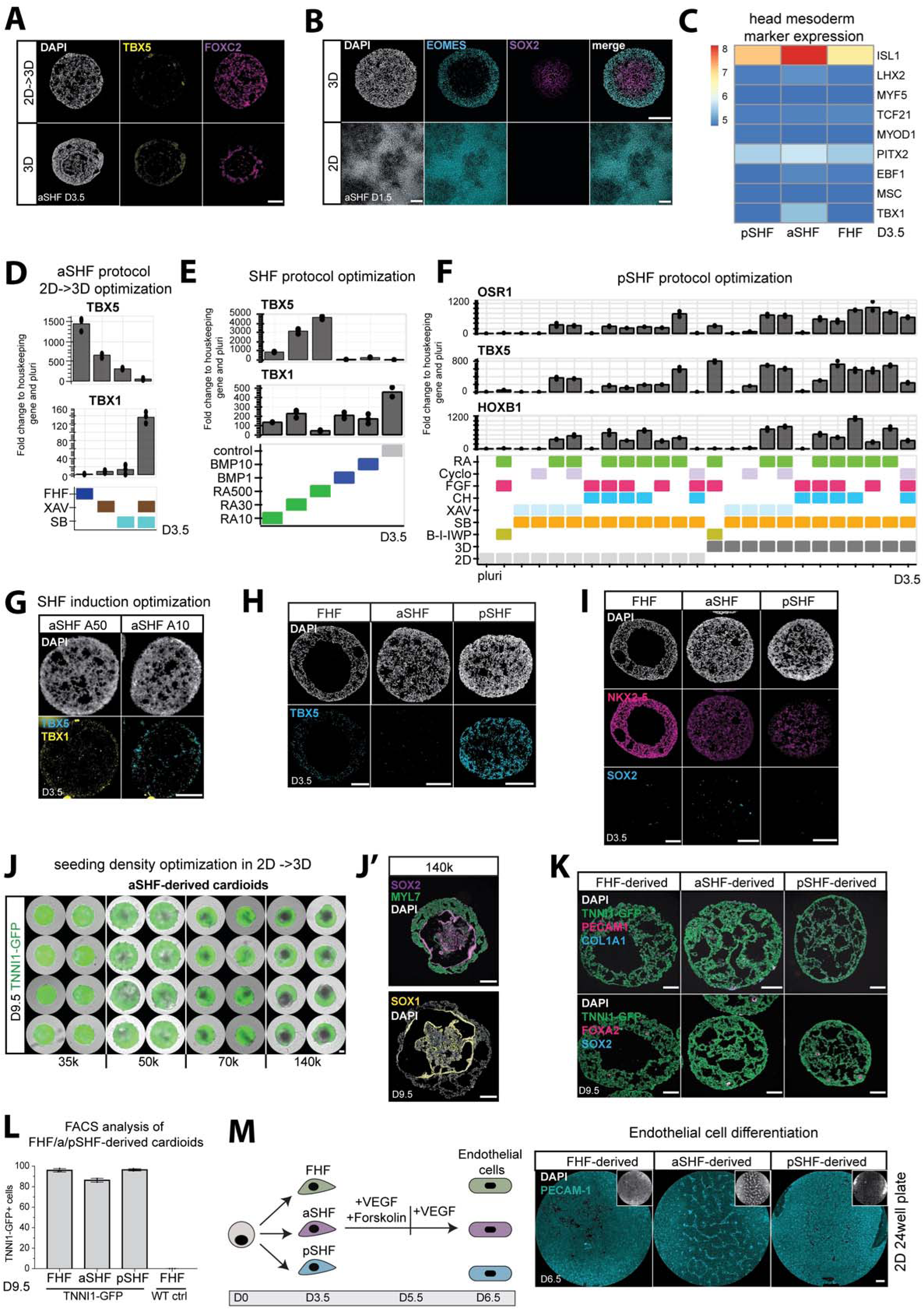
Optimization of the a/pSHF protocol and characterization of a/pSHF-derived cardioids, related to Figure 1. (A) Immunostaining of TBX5 and FOXC2 of aSHF cells at day 3.5 using 3D vs 2D->3D protocol.(B) SOX2 and EOMES staining after SHF mesoderm induction (day 1.5) in 2D, and 3D protocol.(C) Heatmap of bulk RNA-seq analysis at day 3.5 of head mesoderm makers. (D) Optimization of aSHF patterning media 1 (day 1.5 - day 3.5) conditions in 2D -> 3D protocol. RT-qPCR of TBX1 and TBX5 levels at day 3.5. (E) Optimization of a/pSHF protocol by testing different BMP (in ng/ml) and RA concentrations (in nM) during patterning stage 1 using RT-qPCR. (F) Optimization of pSHF protocol by adding different growth factors and small molecules during pSHF patterning stage 1 using 2D and 3D protocols and RT-qPCR. Cyclo: Cyclopamin, CH: CHIR99021, B: BMP4, I: Insulin) (G) RNA-scope staining of TBX1 and TBX5 at day 3.5 of cross-sections of aSHF progenitors induced with different Activin concentrations. (H) TBX5 staining of cross-sections of all three progenitors at d3.5. (I) Immunostaining of NKX2-5 and SOX2 on cross-sections of FHF, aSHF, and pSHF cardioids at day 3.5. (J) Seeding density optimization in 2D-> 3D protocol in aSHF-derived cardioids at day 9.5 derived from TNNI1-GFP reporter line cardioids. (J’) aSHF cardioids started with a high seeding density analyzed with ICC of SOX1/2+ core (neural marker) at day 9.5. (K) Immunostaining of PECAM1 (Endothelial cells), FOXA2 (endoderm), COL1A1 (Fibroblast), and SOX2 (neuroectoderm) in all cardioid subtypes. (L) Quantification of TNNI1-GFP+ cells in FHF, aSHF, and pSHF-derived cardioids at day 9.5 via flow cytometry (N=3, n=8). (M) 2D 24-well plate endothelial cell differentiation of all three progenitor populations. RT-qPCR: Fold change normalized to a housekeeping gene (PBGD) and pluripotency. vst: variance-stabilized transformed counts. All scale bars in this figure have a length of 200µm. Used cell lines in this figure: WTC and H9. All bar graphs show mean ± SD.

**Figure S2:**
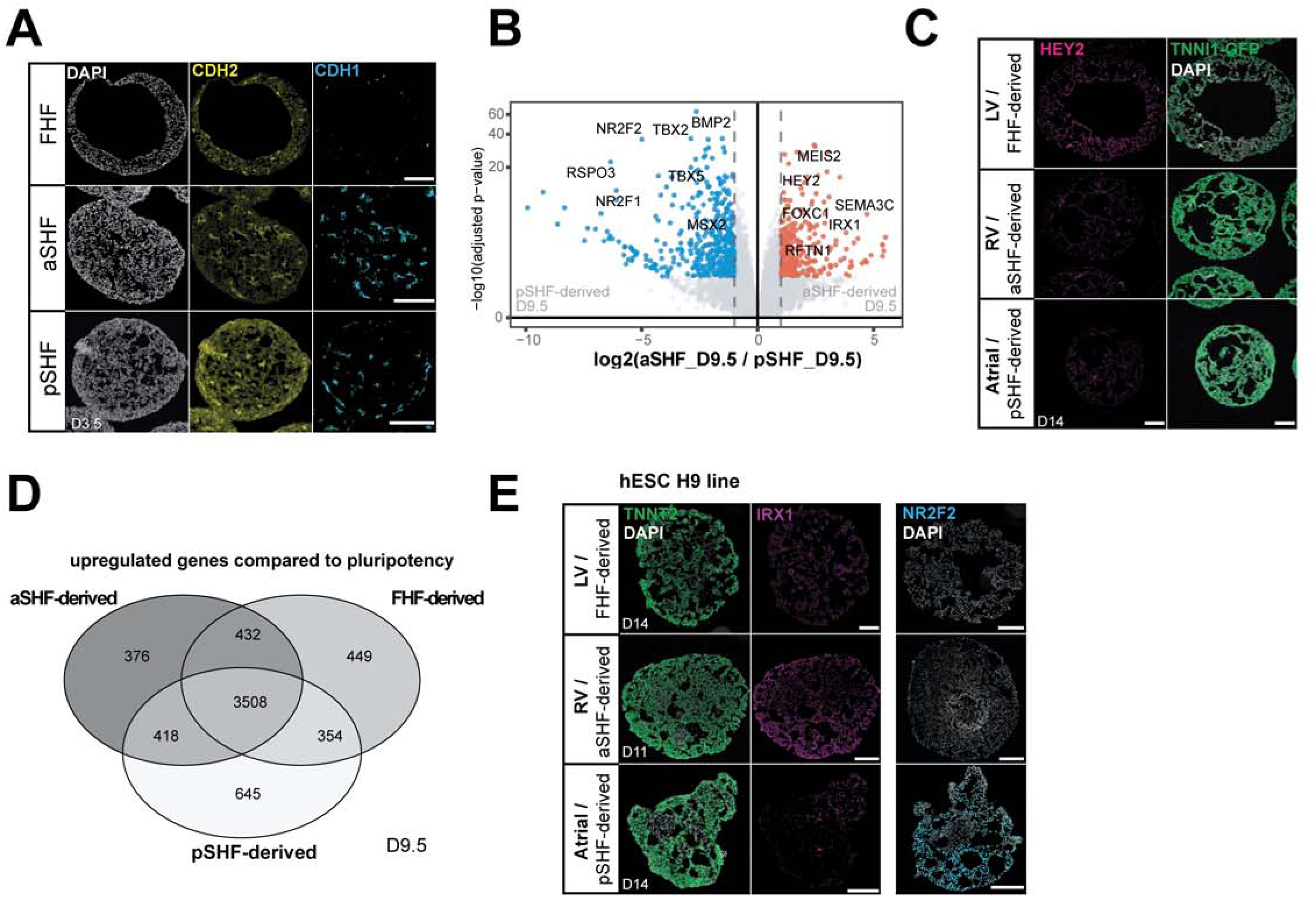
Characterization of FHF-, aSHF- and pSHF-derived cardioids, related to Figure 2. (A) CDH1 and CDH2 immunostaining of FHF, aSHF, and pSHF cardioids at day 3.5. (B) Volcano plot showing the differentially expressed genes at day 9.5 of aSHF- versus pSHF-derived cardioids. (C) Lineage-specific staining of HEY2 (LV marker) in cross-sections of FHF-, aSHF- and pSHF-derived cardioids. (D) Venn Diagram revealing showing the number of upregulated genes compared to pluripotency between FHF-, aSHF- and pSHF-derived cardioids at day 9.5 (E) Immunostaining of Atria (NR2F2) and RV (IRX1) specific genes of cardioids derived from hESC line H9. All scale bars in this figure have a length of 200µm. Used cell lines in this figure: WTC and H9.

**Figure S3:**
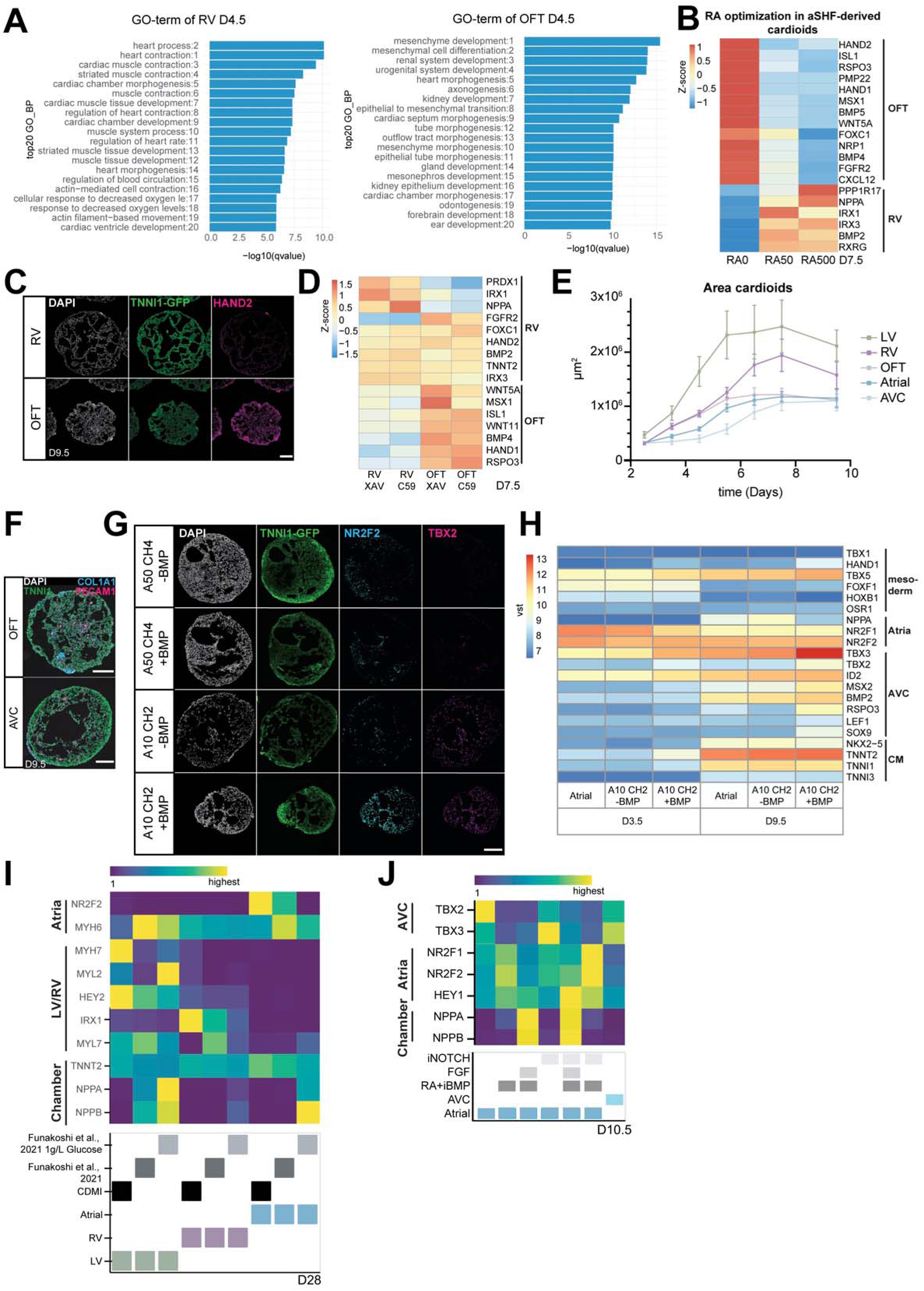
Optimization of OFT and AVC protocol and OFT and AVC cardioid characterization, related to Figure 3. (A) GO-term analysis from bulk RNA-seq upregulated in RV cardioids compared to OFT cardioids at day 4.5 (B) Optimization of RA concentrations during patterning stage 2 using bulk RNA-seq for OFT and RV markers(C) HAND2 staining of OFT and RV cardioid cross-sections at day 9.5. (D) Optimizations of type of iWNT used for RV and OFT markers by bulk RNA-seq (E) Quantification of cardioid area change over time during differentiation in all five protocols (N=3, n=32). (F) COL1A1 (fibroblast marker) and CD31 (endothelial cell marker) staining of OFT and AVC cardioid cross-sections at day 9.5. (G) Optimization of pSHF mesoderm induction conditions using different Activin and CHIR99021 concentration and addition or absence of BMP to distinguish between Atria and AVC cardioids using ICC on cross-sections of NR2F (Atrial marker) and TBX2 (AVC marker). (H) Heat map of bulk RNA-seq for BMP optimization for atrial and AVC genes. (I) RT-qPCR of LV, RV, and atrial cardioids which were treated with published CM chamber specification and maturation conditions (Funakoshi et al., 2021) from day 7.5 to day 28. (I) RT-qPCR of optimization of atrial chamber specification until day 10.5. i: inhibitor, RA: retinoic acid. RT-qPCR: Fold change normalized to a housekeeping gene (PBGD) and pluripotency. vst: variance-stabilized transformed counts. All scale bars in this figure have a length of 200µm.

**Figure S4:**
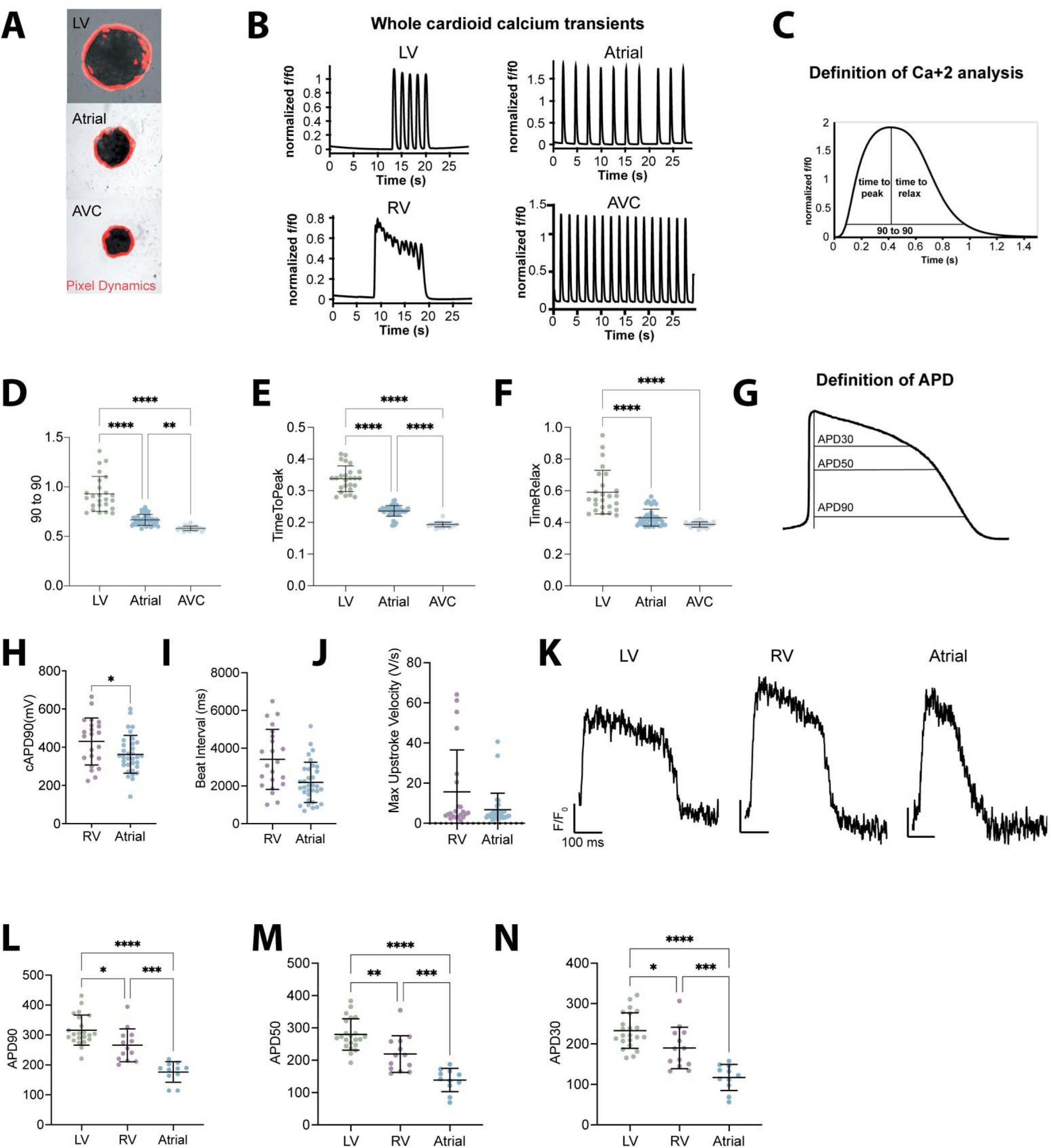
Functional characterization of cardioid subtypes using calcium transients and voltage-sensitive dyes, related to Figure 4. (A) Representative images showing the extent of contraction (red) for different organoid types. (B) Calcium traces showing the relative change in fluorescence intensity (f/f0) were recorded from the whole cardioid area for a time span of 30. (C) Definitions of parameters for subfigures D-F. (D-F) Quantification of whole cardioid Ca^+2^ transients (D) 90 to 90 (E) time to reach max intensity (F) time to relaxation. Data was taken at D9.5. All points represent the mean of each cardioid across all beats recorded. Data was taken from 4 biological replicates with 16 technical replicates per biological replicate for the LV and Atria and 1 biological replicate with 24 technical replicates for the AVC. All cardioids which were not beating were excluded. This resulted in 25, 43, and 27 replicates respectively of the LV, Atria, and AVC. (G) Sketch highlighting how APD’s were calculated for patch-clamp and FluoVolt. (H-I) Additional data for patch-clamp shown in 4I-L. (H) Corrected APD90 values (I) and beat intervals. A correction was performed using the Fridericia formula for QT interval correction. (J) Maximum upstroke velocity. (K) Representative curves from 2D FluoVolt data for LV, RV, atrial cardioids. The y-size bar represents relative intensity change and the x-size bar represents 100 milliseconds (ms). (L-N) APD90, APD50, and APD30 of 2D FluoVolt data. 25 wells for the LV, 26 wells for the LV, and 13 wells for the atria were recorded each point is an average of all APs taken within one well.

**Figure S5.**
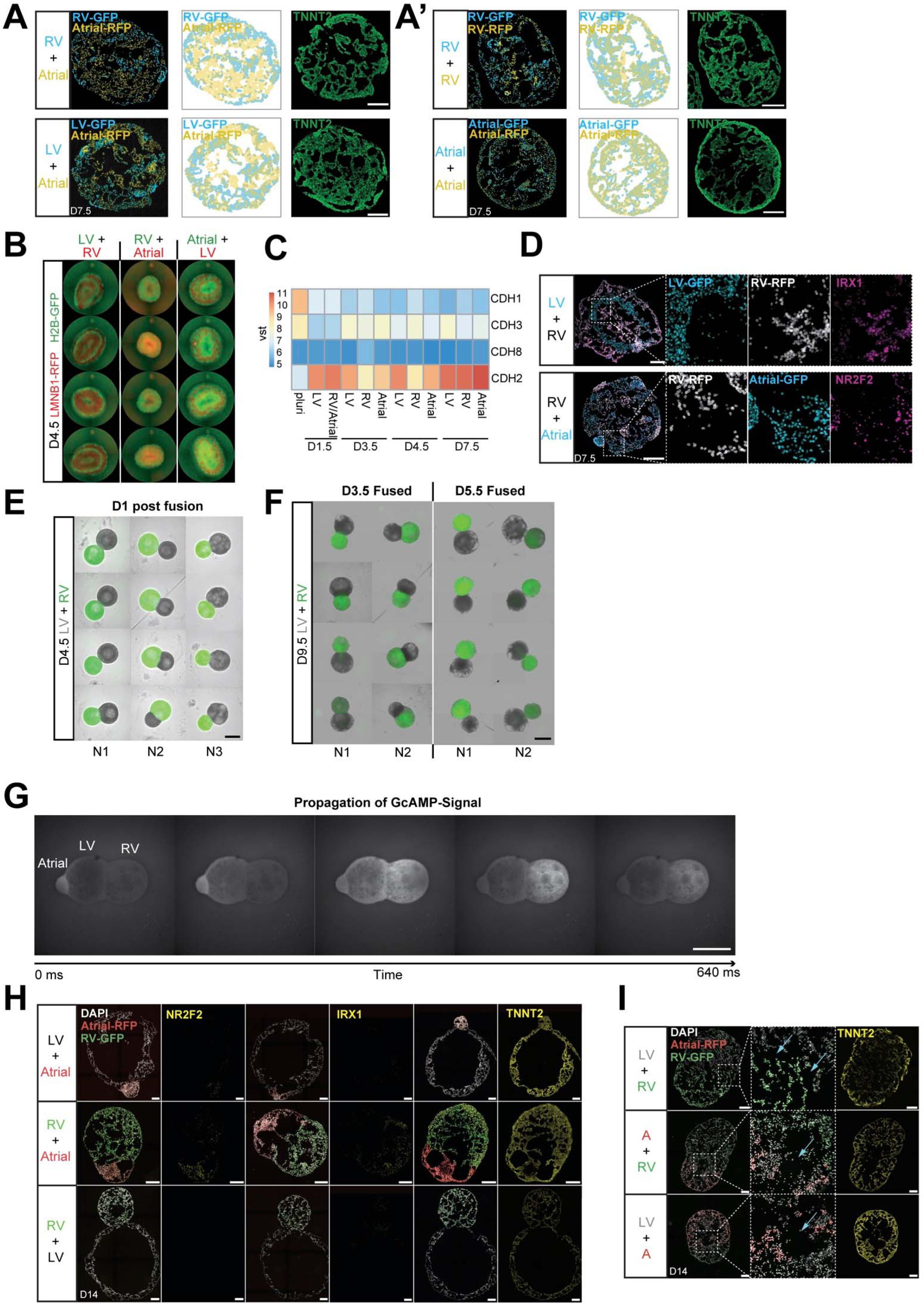
Progenitor sorting and formation of multi-chamber cardioids, related to Figure 5. (A-A’) Cardiac progenitors derived from either H2B-GFP, or LMNB1-RFP hPSC reporter lines were dissociated and mixed at day 3.5. Cross-sections and schematics of cardioids show the sorting of cardiac cells derived from (A) different progenitor populations and (A’) cardioids mixed with the same progenitors. TNNT2 staining shows highly efficient CM differentiation. scale bar, 200 µm. (B) Whole-mount images showing sorting of cardiac progenitors one-day post mixing (day 4.5). (C) Heatmap of bulk-RNAseq analysis of cardioids generated with the normal 2D-3D protocol (non-mixed progenitors) showing differentially expressed Cadherin genes over time. (D) Cross-sections of cardioids mixed with different progenitor populations stained of RV specific (IRX1) and atrial specific (NR2F2) markers. scale bar, 200 µm. (E) Cardioids one day post-fusion (day 4.5). scale bar, 1000 µm. (F) Whole-mount images of cardioids being fused together on day 3.5 or day 5.5. The images were taken on day 9.5. scale bar, 1000 µm. (G) Time lapse of calcium signaling traveling through a three-chambered cardioid. scale bar, 500 µm. (H) Lineage-specific staining (NR2F2 and IRX1) of two-chambered cardioids. scale bar, 200 µm. (I) Cryosection of multi-chambered cardioids of two different compartments on day 14 using protocol depicted in Figure 5M. multi-chambered cardioids share some cavities (indicated by the blue arrow) and express TNNT2. scale bar, 200 µm. vst: variance-stabilized transformed counts.

**Figure S6.**
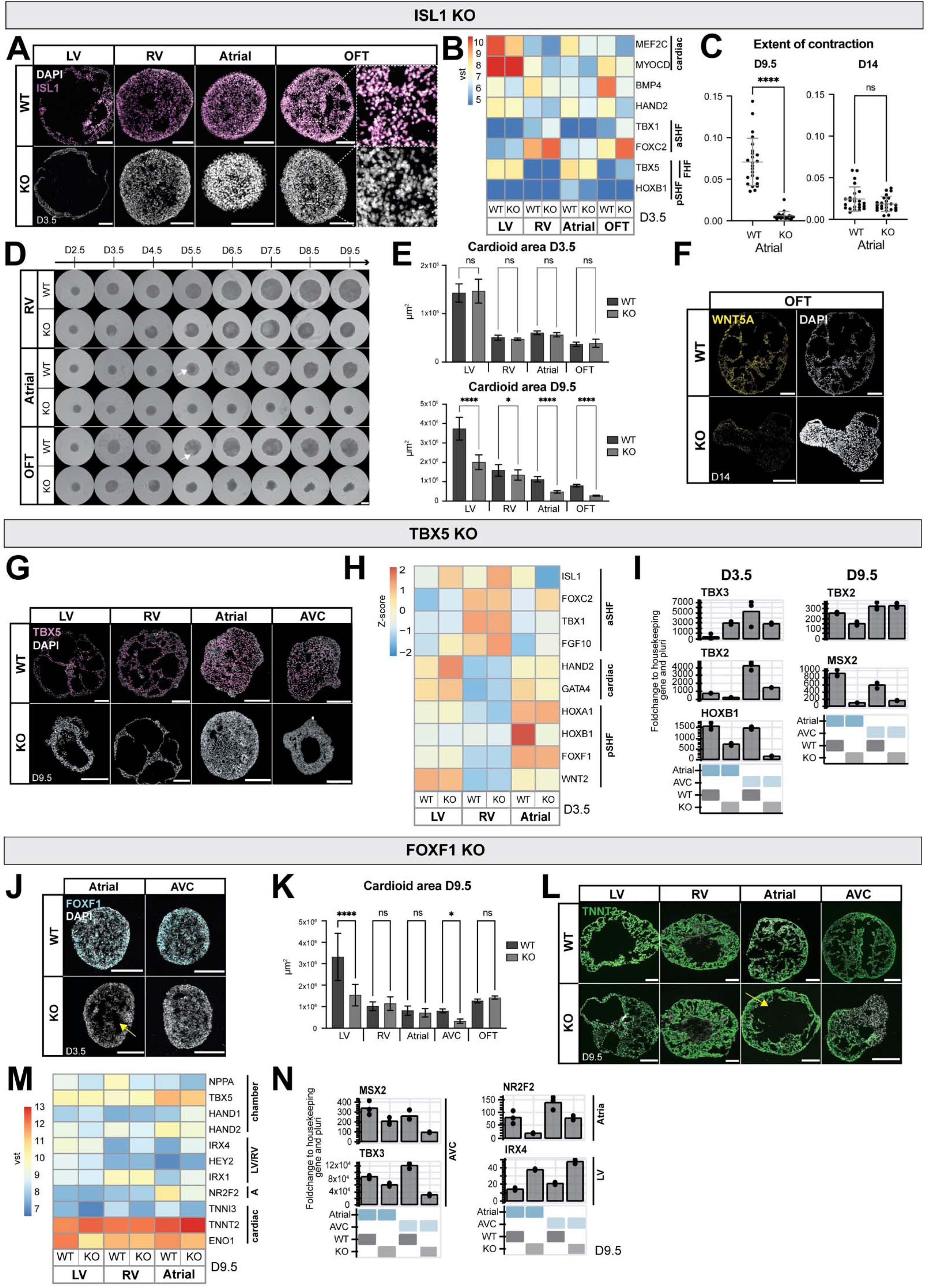
Compartment-specific defects in cardioids with mutations in transcription factors, related to Figure 6. (A) Validation of ISL1 KO line in all protocols at day 3.5 by ICC. (B) Bulk-RNAseq analysis showing misregulated genes in ISL1 KO vs WT at day 3.5. (C) The extent of contraction analysis of atrial ISL1 KO cardioids compared to WT at day 9.5 and day 14 (N=1, n=24). (D) Time course of RV, atrial, and OFT cardioid formation using ISL1 KO and WT line. Arrow indicating cavity formation in WT cardioids. scalebar, 500µm. (E) Quantification of the cardioid area of ISL1 KO and WT cardioids at days 3.5 (N=4, n=8-24) and 9.5 (N=2, n=8-24). (F) Downregulation of WNT5A (OFT marker) in OFT ISL1 KO cardioids compared to WT at day 14 (HCR staining). (G) Validation of TBX5 KO line in LV, RV atrial, and AVC cardioids at day 9.5 by ICC (H) Bulk-RNAseq analysis of TBX5 KO and WT cardioids showing misregulated aSHF and pSHF specific genes at day 3.5. (I) Representative RT-qPCR of atrial and AVC TBX5 KO cardioids compared to WT on days 3.5 and 9.5. F (J) Validation of FOXF1 KO line in atrial and AVC cardioids at day 3.5. Yellow arrow indicating enhanced cavity formation in atrial FOXF1 KO cardioids. (K) FOXF1 KO and WT cardioid area analysis of all cardioid subtypes at day 9.5 (N=3-4 (LV, RV, Atria, AVC) and N=1 (OFT), n=8-16). (L) TNNT2 expression in FOXF1 WT vs KO cardioids at day 9.5. Yellow arrow indicating increased cavity in FOXF1 KO atrial cardioids compared to WT. (M) Bulk-RNAseq analysis showing misregulated genes of LV, RV, and atrial cardioids using FOXF1 KO line compared to WT line at day 9.5. (N) Representative RT-qPCR of atrial and AVC cardioids in FOXF1 KO line compared to WT line. All scale bars in this figure have a length of 200µm. RT-qPCR: fold change normalized to a housekeeping gene (PBGD) and pluripotency. vst: variance-stabilized transformed counts. All bar graphs show mean ± SD. For all statistics, one-way ANOVA was used. *p < 0.05, **p < 0.01, *** p < 0.001, ****p < 0.0001. ns: not significant.

**Figure S7.**
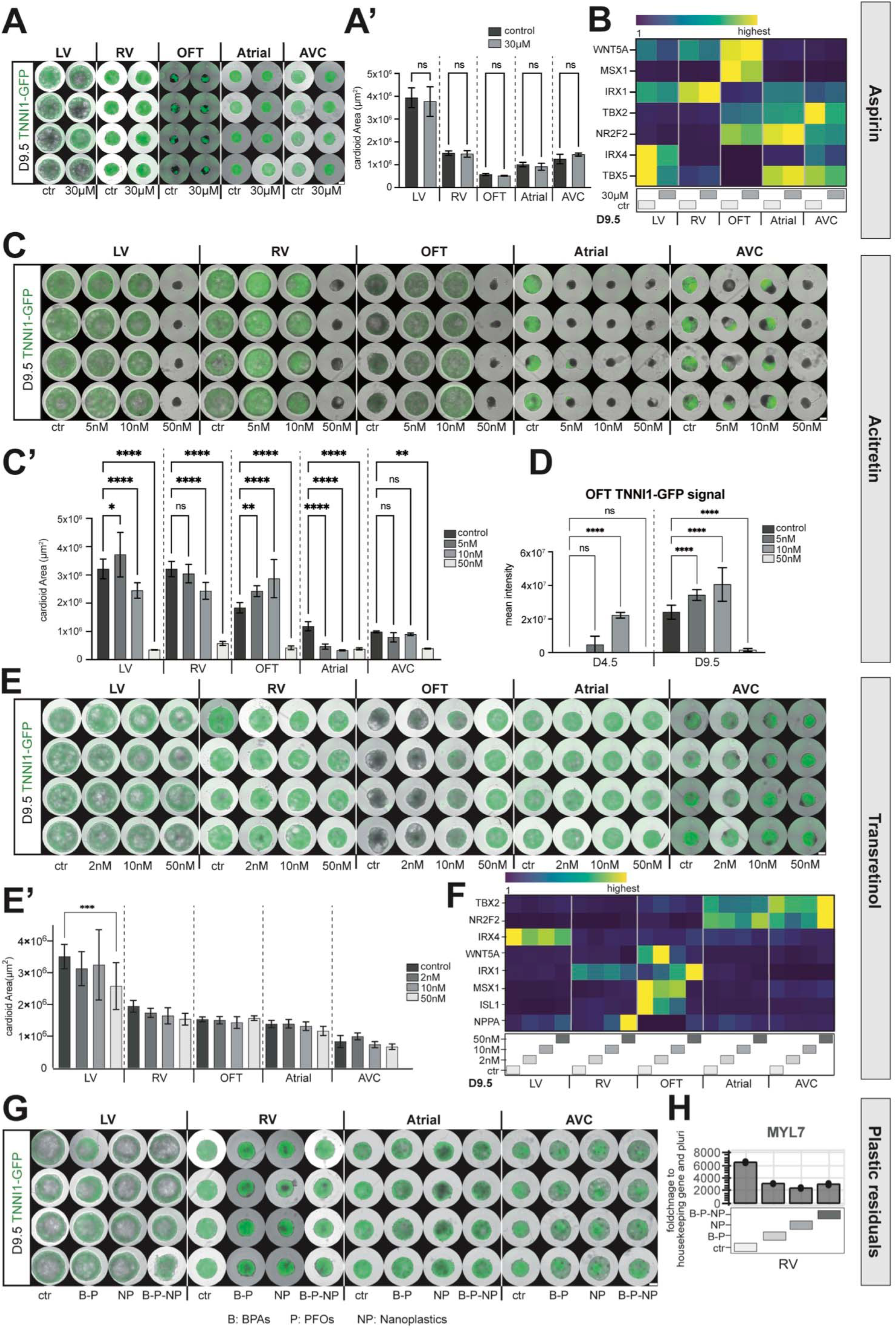
Characterization of teratogen-induced compartment-specific defects in cardioids, related to Figure S7. (A) Whole-mount images and (A’) quantification of area of cardioids derived from TNNI1-GFP reporter line treated with Aspirin compared to untreated cardioids. day 9.5 (N=1, n=8). Ctr: control (B) RT-qPCR of cardioids treated with Aspirin compared to control cardioids. (C) Representative whole-mount images of cardioids treated with different concentrations of acitretin and (C’) quantification of the cardioid area at day 9.5 (N=1, n=8). (D) TNNI1-GFP reporter signal qualification in OFT cardioids treated with acitretin compared to untreated cardioids at day 4.5 and 9.5. (N=1, n=8) (E) Whole-mount images and quantification (E’) of cardioids derived from TNNI1-GFP reporter line treated with trans-retinol at day 9.5 (N=1, n=8). (F) RT-qPCR shows misregulated genes of all cardioid subtypes induced with trans-retinol. (G) Whole-mount images of cardioids derived from TNNI1-GFP reporter line treated with different combinations of Plastic residuals. (N=1, n=8). (H) Representative RT-qPCR showing MYL7 expression in RV cardioids induced with plastic residuals compared to control RV cardioids. All cardioids were treated with teratogens from mesoderm induction (day 0) until day 9.5. All scale bars in this figure have a length of 500µm except where specified. RT-qPCR: Fold change normalized to a housekeeping gene (PBGD) and pluripotency. All bar graphs show mean ± SD. For all statistics, one-way ANOVA was used. *p < 0.05, **p < 0.01, *** p < 0.001, ****p < 0.0001. ns: not significant.

## SUPPLEMENTARY INFORMATION TITLES

Video S1: **Cardioid showing different origins of signal propagation.** Video shows calcium transients across an atrial cardioid at day 9.5. Video is slowed down to one-tenth the speed for ease of viewing the spot of signal propagation.

Video S2: **Two-chambered cardioid contracting in a coordinated manner.** A LV cardioid (right) paces a RV cardioid (left). The phase-contrast video was recorded at 100 frames per second.

**Video S3: 3-chambered cardioid showing the direction of signal propagation**

Video shows calcium transients in an atria-LV-RV co-developed cardioid at D9.5. The video was recorded at 50 frames per second.

**Table S1:**
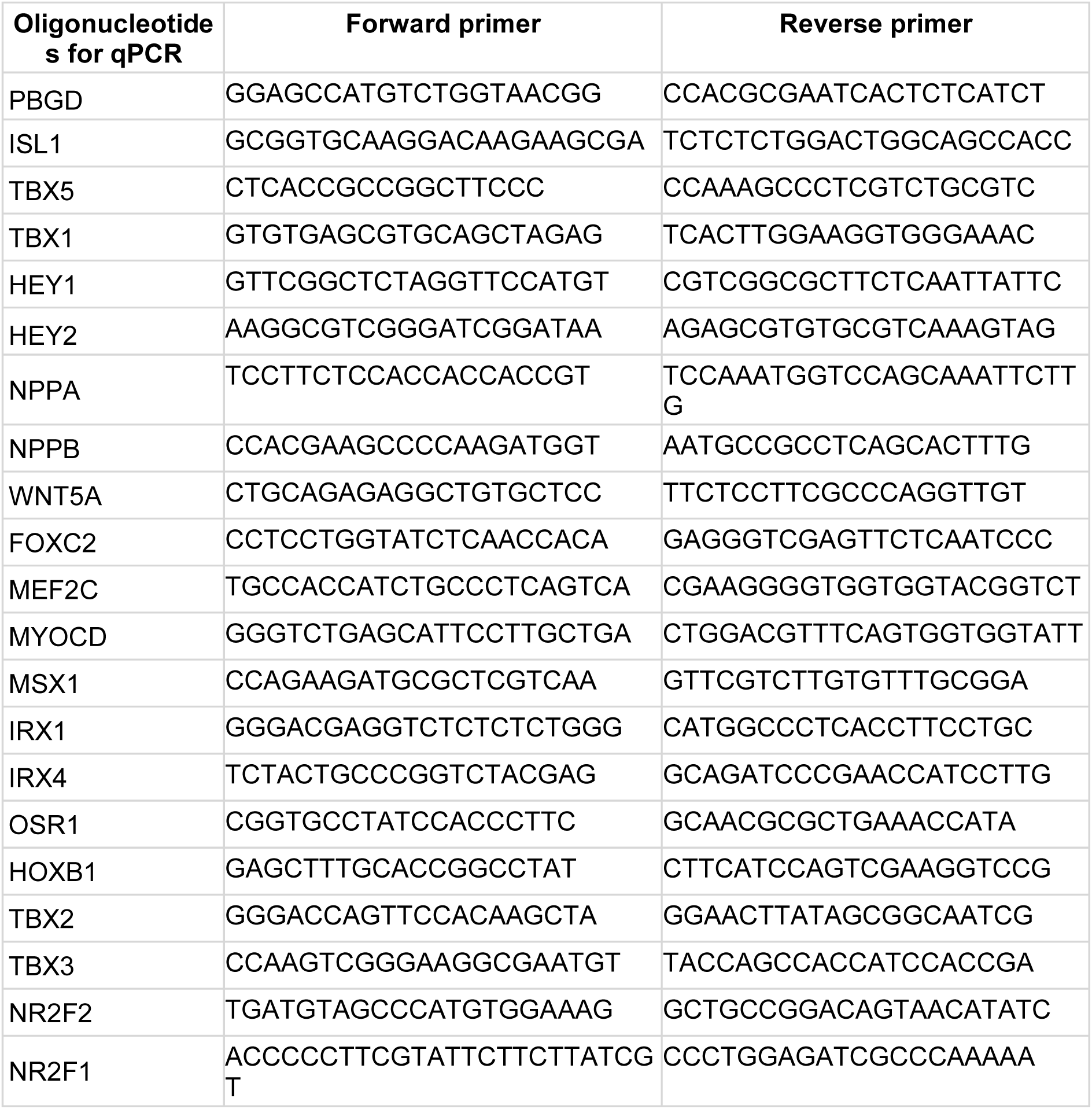
Oligos used for RT-qPCRS.

**Table S2:** Maximum fold-change for RT-qPCR results.

| Gene | Max Foldchange |
| --- | --- |
| Figure 3L |  |
| MYL2 | 5454 |
| MYH6 | 1510 |
| MYH7 | 36 |
| NPPB | 95268 |
| NPPA | 21982 |
| IRX4 | 1039 |
| IRX1 | 169 |
| NR2F2 | 147370 |
| TBX3 | 326 |
| TBX2 | 9 |
| Figure S3I |  |
| NRF2F2 | 133 |
| MYH7 | 5471 |
| MYH6 | 515323 |
| MYL2 | 45559 |
| HEY2 | 35 |
| IRX1 | 761 |
| MYL7 | 2054 |
| cTNT | 21347 |
| hTTNT2 | 2764 |
| NPPA | 4216399 |
| NPPB | 30693 |
| Figure S3J |  |
| TBX2 | 16004 |
| TBX3 | 33003 |
| NR2F1 | 4778 |
| NR2F2 | 14379 |
| HEY1 | 88 |
| NPPA | 1227082 |
| NPPB | 260 |
| Figure 7B |  |
| IRX4 | 150 |
| NPPA | 11570 |
| TBX2 | 4083 |
| Figure 7B cont. |  |
| NR2F2 | 5802 |
| IRX1 | 112 |
| WNT5A | 150 |
| Figure 7E |  |
| TBX5 | 15653 |
| FOXF1 | 1614 |
| HOXB1 | 138869 |
| TBX1 | 479 |
| OSR1 | 256 |
| FOXC2 | 4613 |
| MEF2C | 1505 |
| MYOCD | 1248 |
| WNT5A | 162 |
| NPPA | 2148265 |
| TBX2 | 75090 |
| IRX4 | 239 |
| IRX1 | 11 |
| Figure S7B |  |
| WNT5A | 205 |
| MSX1 | 303 |
| IRX1 | 21 |
| TBX2 | 31494 |
| NR2F2 | 704 |
| IRX4 | 712 |
| TBX5 | 31792 |
| Figure S7F |  |
| TBX2 | 13675 |
| IRX1 | 267 |
| NR2F2 | 11757 |
| IRX4 | 920 |
| MSX1 | 549 |
| WNT5A | 199 |
| ISL1 | 4750 |
| NPPA | 596358 |

**Table S3:** Reason for cardioid exclusion for Ca^+2^ signal propagation analysis. *numbers represent the number of cardioids excluded in each category

|  | D6.5 |  |  |  | D9.5 |  |  |  |
| --- | --- | --- | --- | --- | --- | --- | --- | --- |
|  | LV | RV | Atria | AVC | LV | RV | Atria | AVC |
| no beats | 4 | 37 | 8 | 3 | 14 | 43 | 1 | 5 |
| no directionaliy | 13 | 2 | 5 | 7 | 4 | 0 | 5 | 5 |
| 2 origins | 2 | 0 | 0 | 0 | 0 | 0 | 1 | 0 |
| small area | 1 | 0 | 1 | 5 | 0 | 0 | 1 | 0 |

